# Sequence-encoded conformational biases shape self-assembly modes of intrinsically disordered proteins

**DOI:** 10.1101/2025.10.27.683987

**Authors:** Ryoga Kobayashi, Norio Yoshida, Yohei Miyanoiri, Hideki Nakamura, Miu Ekari, Emi Sakamoto, Misato Tsutsumi, Kayo Imamura, Takashi S. Kodama, Hidehito Tochio, Naotaka Sekiyama

## Abstract

Self-assembly of intrinsically disordered proteins (IDPs) underlies cellular functions and disease pathogenesis. This process is mediated by two intermolecular interaction modes: transient point-to-point contacts described by the sticker-and-spacer framework, and persistent surface-to-surface contacts proposed in the cross-β hypothesis. We investigated the molecular basis of these modes in the context of conformational biases, defined as sequence-encoded structural preferences of local segments. Using a five-residue model, we generated lag-series IDPs from the TIA-1 prion-like domain by systematically modulating conformational biases while preserving amino acid composition. The lag-series IDPs demonstrated distinct condensate properties and varying capacities for amyloid fibril formation. Their structural analyses revealed that strongly biased regions preferentially adopt extended structures, including β-strands, and the spacing between these regions influences metastable β-sheet formation. Our findings demonstrate that local conformational biases shape interaction modes of IDPs, thereby linking sequence to condensate properties and amyloid fibril formation.

**Significance Statement:** Proteins fold into three-dimensional structures defined by their amino acid sequences. In contrast, intrinsically disordered proteins (IDPs) lack stable structures, yet their sequences encode unique self-assembly behaviors, including phase separation and amyloid fibril formation. Can such behaviors be explained within structure-based frameworks? Using a five-residue model, we showed that IDP sequences encode local structural preferences, termed conformational biases, within short segments. These biases determine whether segments engage in transient point-to-point interactions driving phase separation or persistent surface-to-surface interactions leading to amyloid fibril formation. By bridging sequence, structure, and interaction modes, our work provides a comprehensive mechanism for self-assembly and conceptual tools for understanding IDP-related biological functions and disease mechanisms.

## Introduction

Intrinsically disordered proteins (IDPs) are abundant in eukaryotic proteomes and contribute to various cellular functions, including transcriptional regulation, cell signaling, and the assembly of membraneless organelles (MLOs) (*1, 2*). Unlike folded proteins, IDPs do not adopt a single, well-defined structure; instead, they populate highly dynamic conformational ensembles. Within these ensembles, short sequence segments, typically several residues, exhibit sequence-encoded structural preferences or conformational biases (*3, 4*). These conformational biases enable unique protein-protein interactions through diverse binding modes, such as induced folding, conformational selection, and fuzzy complexes (*5–8*). Elucidating how the amino acid sequence encodes such conformational biases is critical for decoding the structure-function relationships in IDPs.

Many IDPs localized in MLOs have been reported to undergo phase separation through self-assembly, forming condensates that exhibit diverse physical properties, ranging from liquid-like droplets to solid-like gels (*9*). As dynamic hubs, MLOs regulate cellular processes by mediating reversible interactions among defined sets of biomolecules. In some cases, these condensates convert into highly ordered filamentous protein aggregates, amyloid fibrils (*10–13*). Amyloid fibrils were initially identified as pathological deposits associated with neurodegenerative diseases, such as Alzheimer’s and Parkinson’s diseases (*14*), yet recent studies have revealed physiological roles, such as long-term memory and signal transduction, across diverse organisms (*15, 16*). Despite the broad involvement of IDP self-assembly in cellular functions and disease pathogenesis, the underlying molecular mechanisms remain largely elusive.

Structural studies have revealed that IDPs retain largely disordered conformations within condensates and engage in multivalent, residue-level interactions, including electrostatic, hydrophobic, π-π, π-cation, π-sp^2^, and backbone hydrogen bonding (*17– 20*). These findings led to the development of the sticker-and-spacer framework, which represents an IDP as a beads-on-a-string polymer whose residues act either as cohesive stickers or flexible spacers based on its physicochemical properties (*19, 21*). By imposing constraints between stickers while allowing the beads to move randomly, this model can capture key phase separation properties, such as transition temperature, saturation concentration, and viscosity (*19, 22–25*). Furthermore, recent studies based on this framework have shown that IDPs adopt preferential orientations and compact conformations at the surface of condensates (*23, 26*), suggesting that these surfaces may act as hotspots for amyloid fibril formation. Together, the transient, residue-level interactions and conformational dynamics appear to govern IDP self-assembly.

An alternative hypothesis of IDP self-assembly mechanism, hereafter the cross-β hypothesis, proposes that certain IDPs form condensates through transient cross-β interactions. A cross-β architecture, the hallmark of amyloid fibrils, consists of β-strands oriented perpendicular to the fibril axis and stabilized by interstrand hydrogen bonds (*27*). Short sequence motifs of 6–10 residues that can adopt such cross-β registers have been identified in many self-assembling IDPs (*28–33*). Inside condensates, these motifs are suggested to transiently sample cross-β interactions (*34– 36*), and when the interactions persist long enough to cross a nucleation threshold, they can trigger and propagate amyloid fibril formation. Structural studies of these short motifs have revealed labile architectures, such as kinked backbones (*30, 31*), extended β-strands (*32, 33*), and stacks of charged residues (*29*), consistent with reversible assembly of IDP condensates. While these observations support the cross-β hypothesis, solution NMR and Raman spectroscopy studies detect little or no ordered structures inside condensates (*18, 37*), indicating that debate over the hypothesis remains.

Taken together, two models have been proposed to explain IDP self-assembly. The sticker-and-spacer framework treats an IDP as a beads-on-a-string polymer whose residues sample random conformations and form transient, residue-level contacts. We refer to these as point-to-point interactions. By contrast, the cross-β hypothesis proposes that short sequence segments engage in more persistent contacts that align to form cross-β architectures. We refer to these as surface-to-surface interactions. During IDP self-assembly, both interaction modes can act in concert: some regions of an IDP may be dominated by point-to-point interactions, whereas others may be characterized by surface-to-surface interactions. However, the molecular factors that determine which type of interaction mode predominates for a given sequence remain unclear.

In this study, we investigated the molecular basis of these modes in the context of conformational biases, defined as sequence-encoded structural preferences of local segments. To this end, we employed a five-residue model (F-r model), previously derived from NMR relaxation data of the T-cell intracellular antigen-1 prion-like domain (TIA-1 PLD) (*38, 39*). The F-r model yields an F-r score for each residue, with higher scores indicating stronger conformational biases and lower scores indicating weaker conformational biases. By systematically modulating the periodicity of F-r scores while preserving amino acid composition, we designed artificial IDPs derived from the TIA-1 PLD sequence, hereafter referred to as lag-series IDPs. The lag-series IDPs exhibited distinct condensate properties and a propensity of amyloid fibril formation. To investigate the underlying mechanisms, we performed structural studies using all-atom molecular dynamics (MD) simulations, nuclear magnetic resonance (NMR), circular dichroism (CD) spectroscopy, and AlphaFold2-based structure prediction. Our results suggest that high F-r score segments preferentially adopt extended structures, including β-strands, that enable surface-to-surface interactions. Moreover, the spacing between high F-r score segments dictates whether those regions form metastable β-sheets, which we consider as hotspots for amyloid nucleation. Our results show that sequence-encoded conformational biases shape the balance of interaction modes in IDPs, thereby linking sequence to condensate properties and amyloid fibril formation.

## Results

### Designing lag-series IDPs with periodic conformational biases

In this study, we hypothesize that the choice between point-to-point and surface-to-surface interactions is dictated by conformational biases of short segments within the amino acid sequence (Fig. 1A). Here, conformational bias denotes a sequence-encoded structural preference of a short segment, typically several residues (e.g., five residues), for particular backbone geometries. Segments with high backbone flexibility sample a broad conformational space and exhibit fast dynamics (see Fig. 1A, segment depicted as “weak bias”), while segments with restricted flexibility are confined to a narrow conformational space and exhibit slow dynamics (see Fig. 1A, segment depicted as “strong bias”). We propose that point-to-point interactions predominate in weakly-biased segments, since they are less likely to form persistent surfaces, making transient residue-level contacts more favorable. Conversely, surface-to-surface interactions predominate in strongly-biased segments, due to persistent contacts between multiple residues.

**Fig. 1.**
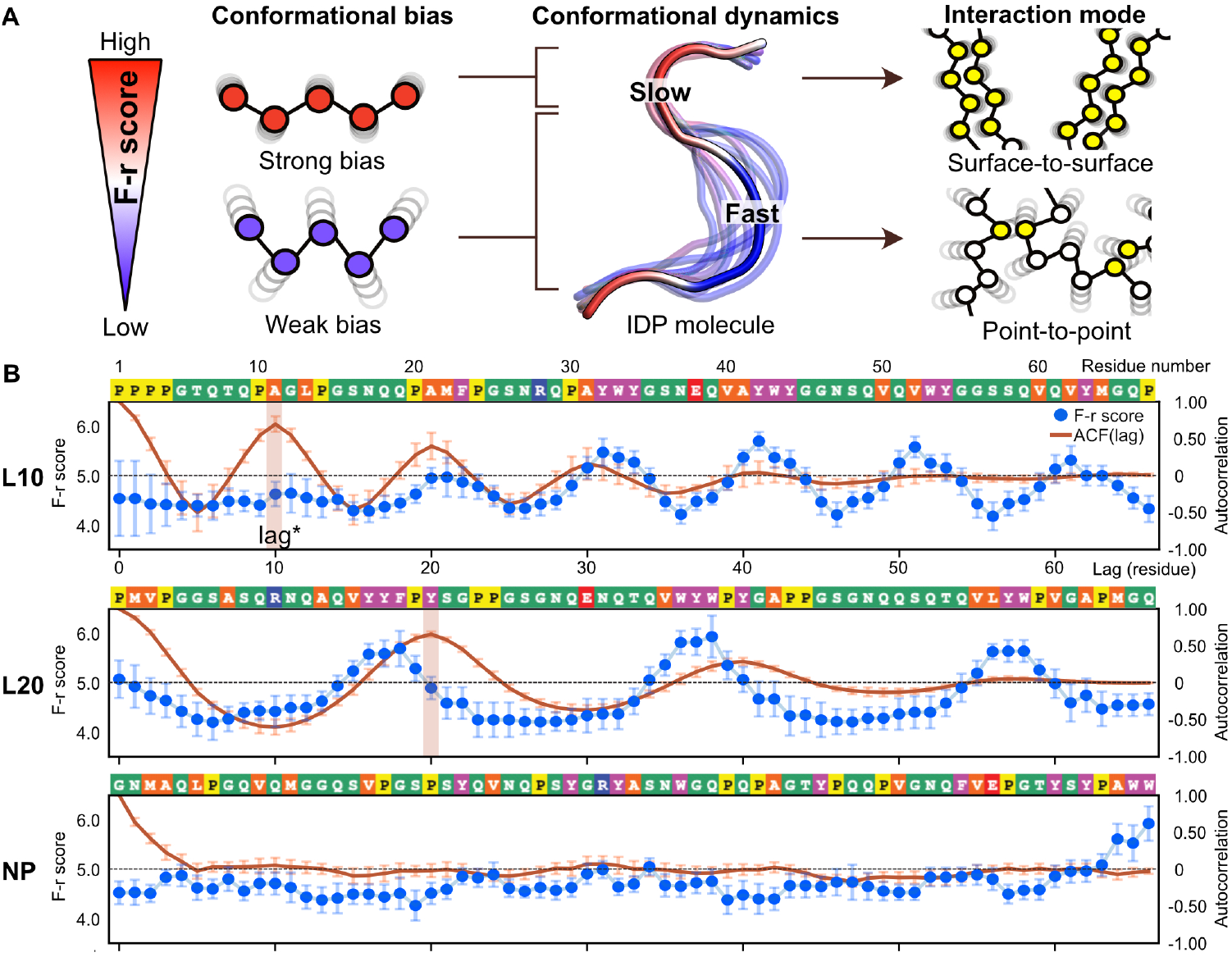
Conformational biases dictate interaction modes of IDPs. **(A)** Schematic diagram of our hypothesis: F-r scores quantify local conformational biases within IDP segments, with high F-r scores indicating strong conformational biases and low F-r scores indicating weak conformational biases. We propose that segments with weak conformational biases exhibit fast dynamics and favor point-to-point interactions, while segments with strong conformational biases exhibit slow dynamics and favor surface-to-surface interactions. **(B)** F-r scores and autocorrelation functions ACF(lag) of lag-series IDPs. L30, L40, L50, and WT are shown in Fig. S2. Each panel shows the F-r scores and autocorrelation functions ACF(lag). F-r scores are calculated from 50 different physicochemical properties of amino acids. Blue data points represent the mean value of the F-r scores, with error bars indicating the standard deviation (SD) of F-r scores. Red solid lines represent the mean value of ACF(lag) calculated from 50 types of F-r scores, with error bars indicating the SD of the ACF(lag) values. Amino acid sequences are shown above the graphs, and colors for each residue indicate the amino acid types: green for polar (G, T, Q, S, N), orange for hydrophobic (A, L, M, V), magenta for aromatic (F, W, Y), yellow for proline (P), red for acidic (E), and blue for basic (R) residues.

To test this hypothesis, we applied a five-residue model (F-r model), originally derived from the structural analysis of TIA-1 PLD (*38*). This model converts amino acid sequences into indices representing physicochemical properties. The geometric mean of these values over consecutive five residues provides an estimate of relaxation rates. We refer to these values as five-residue scores (F-r scores). While examining this model, we conceived that the F-r score could serve as an indicator of conformational biases in local segments. This idea is supported by previous studies combining NMR and MD simulations, which demonstrated that IDP dynamics, reflected in relaxation rates, are predominantly governed by the coordinated motions of short segments, particularly those forming secondary structures (*40–43*). These findings led us to reason that IDP dynamics emerge from innate conformational biases of short segments encoded in the sequence. In this framework, higher F-r scores represent stronger conformational biases, whereas lower scores reflect weaker ones. According to this hypothesis, the distribution of F-r scores should provide a quantitative framework linking conformational biases to IDP dynamics and interaction modes.

Building on this rationale, we hypothesized that tuning the periodic arrangement of conformational biases along an IDP sequence could shift the balance between point-to-point and surface-to-surface interactions, thereby altering condensate properties. This approach was inspired by previous studies showing that the distribution of sticker residues, such as aromatic or charged residues, significantly impacts condensate properties (*19, 22, 24*). To systematically modulate the periodicity of conformational biases in the TIA-1 PLD sequence, hereafter wild-type (WT), we generated artificial IDP sequences by shuffling the WT sequence and selecting variants with strong periodicity of F-r scores. Periodicity was quantified by the autocorrelation function ACF(lag), where lag^*^ represents the spacing between high F-r score regions (Fig. S1). This procedure yielded artificial sequences named L10, L20, L30, L40, and L50, corresponding to their respective lag^*^ values. Each sequence exhibited a clear peak in the ACF(lag) at the intended lag value, confirming their expected periodicity (Figs. 1B, S2 and S3A). To generate a sequence with low periodicity (non-periodic sequence, NP), we selected the sequence with minimal summed ACF(lag). NP had an average ACF(lag) of -0.02, confirming the absence of periodicity. We refer to these artificial IDPs as lag-series IDPs.

Among these sequences, we observed that the lengths of contiguous regions with high F-r scores (greater than 5.0), which we refer to as high F-r score regions, increased with longer lag (Fig. S3B). High F-r score regions were enriched in hydrophobic and aromatic residues, while low F-r score regions were enriched in polar residues (Figs. 1B and S2). By tuning the F-r scores, we could systematically control both the length of biased regions and the spacing between them.

### Condensate properties of lag-series IDPs

To examine the condensate properties of lag-series IDPs, we prepared recombinant proteins of each sequence and diluted the purified proteins in a neutral pH buffer. Among these, L20, L40, L50, and NP formed spherical phase-separated condensates, similar to those of WT (Fig. 2A). In contrast, L30 formed condensates composed of small particles, while L10 produced amorphous aggregates. To assess the temperature-dependent reversibility of these condensates, we heated these condensates at 95 °C for 10 min. All condensates, except for L10, fully dissolved at this temperature, as indicated by the transparency of the test tubes. Notably, the L10 aggregates remained intact, indicating thermal irreversibility (Fig. 2B).

**Fig. 2.**
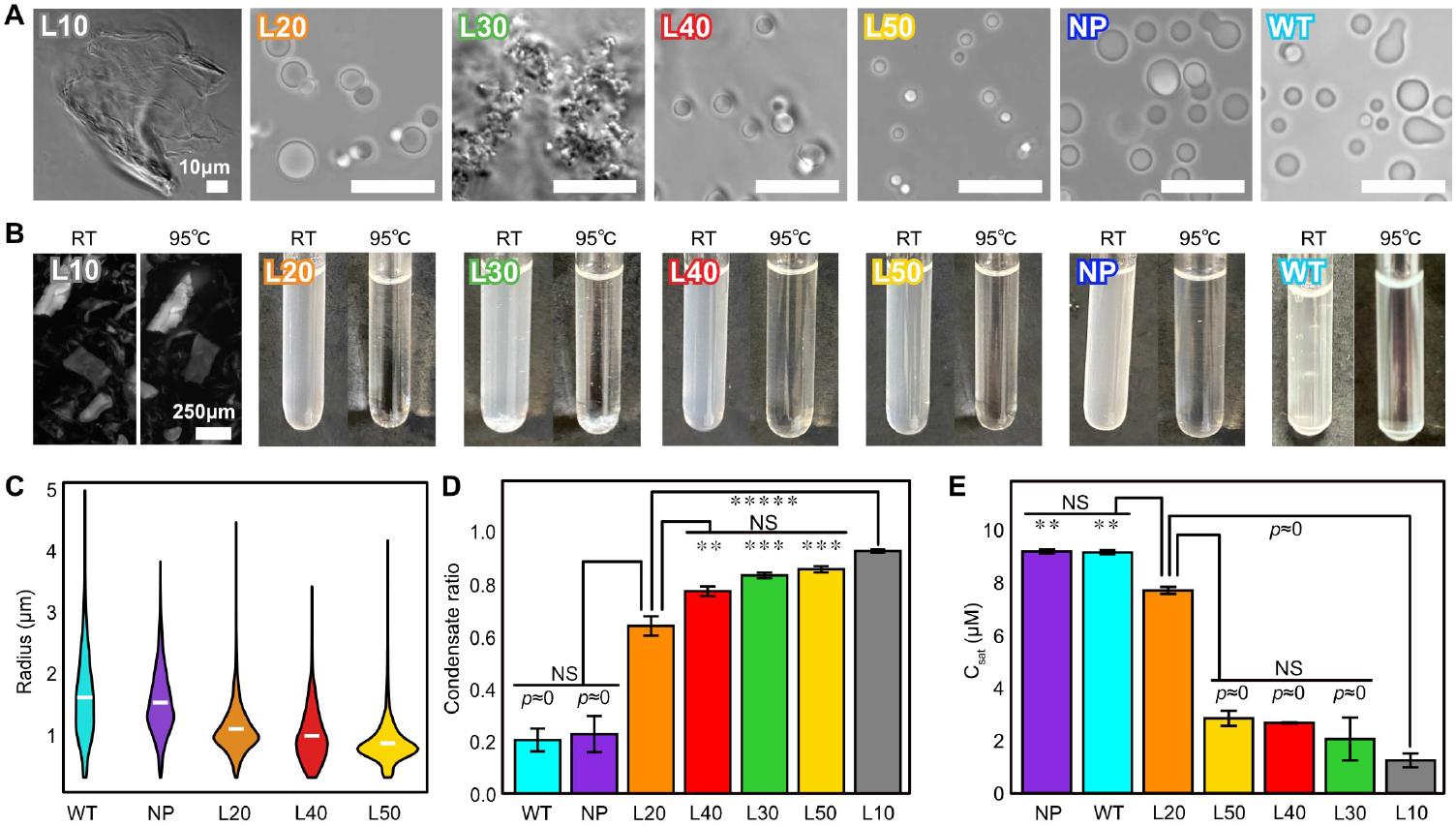
Condensate properties of lag-series IDPs. **(A)** Bright-field microscopy images of lag-series IDP condensates. Scale bar, 10 μm. **(B)** Temperature-dependent reversibility. Left and right panels show room temperature (RT) and 95 ℃, respectively. For L10, bright-field images are shown due to minimal visible change in tubes. Scale bar, 250 µm. **(C)** Condensate size distributions. Radii were measured from the fluorescence images (Fig. S4A). White markers in violin plots indicate mean; edges show maximum/minimum. N (particles) = 3407, 3370, 4653, 1350, and 2394 for WT, NP, L20, L40, and L50, respectively. All pairs significantly different (*p* < 1.0 × 10^−7^) via Steel-Dwass test. **(D)** Condensate ratio in 8 % 1,6-HD buffer. Bar plots represent mean ± SD (N = 3). Tukey’s method was used for multiple comparisons. Asterisks represents adjusted *p*-value vs. L20, ^**^: *p* < 5.0 × 10^-3, ***^: *p* < 5.0 × 10^-4, *****^: *p* < 5.0 × 10^-6^, *p* ≈ 0: *p* < 1.0 × 10^-7^, and NS: no significant difference. **(E)** Saturation concentrations (C_sat_). Bar plots represent mean ± SD (N = 3). Tukey’s method was used for multiple comparisons. Asterisks represents adjusted *p*-value vs. L20, ^**^: *p* < 5.0 × 10^-3^, *p* ≈ 0: *p* < 1.0 × 10^-7^, and NS: no significant difference..

Among the spherical condensate-forming lag-series IDPs (L20, L40, L50, NP, and WT), we observed differences in condensate sizes (Fig. S4A). Microscopic image analysis revealed that NP and WT formed the largest condensates with broad size distribution, followed by L20, L40, and L50 in descending order of size (Fig. 2C). We surmise that larger and more diverse condensate sizes reflect higher fluidity due to frequent fusion and fission events, whereas smaller and more uniform sizes suggest lower fluidity. These results suggest that NP and WT form more fluid condensates than other lag-series IDPs.

To evaluate solubility of lag-series IDPs, we measured the condensate ratio in 1,6-hexanediol (1,6-HD) using tryptophan fluorescence. The condensate ratio is the proportion of IDP remaining in condensates relative to the total amount of IDPs. A higher condensate ratio under high 1,6-HD concentrations indicates lower solubility. All condensates, except for L10, dissolved in a concentration-dependent manner with 1,6-HD (Fig. S4B). L10, however, remained insoluble even at 16 % 1,6-HD concentration. At 8 % 1,6-HD, statistical analysis revealed that WT and NP had the highest solubility, followed by L20 with intermediate solubility, while L30, L40, and L50 exhibited lower solubility (Fig. 2D).

We also determined the saturation concentration (C_sat_) in 8 % 1,6-HD. Consistent with the condensate ratio results, WT and NP showed the highest C_sat_ values (> 9 μM), followed by L20 (7.65 µM). L30, L40, and L50 exhibited lower C_sat_ values (2-3 μM), and L10 demonstrated the lowest (1.24 μM) (Fig. 2E).

In summary, these results suggest that lag-series IDPs exhibit distinct condensate properties depending on their periodicity: NP, like WT, forms fluid and soluble condensates; L20 shares this tendency to a lesser degree, whereas L30, L40, and L50 tend to form less fluid and less soluble condensates, and L10 uniquely forms irreversible aggregates.

### Amyloid fibril formation of lag-series IDPs

We next asked whether condensates formed by lag-series IDPs mature into amyloid fibrils. To monitor amyloid fibril formation, we performed the thioflavin-T (ThT) assay, in which the fluorescence of ThT increases upon binding to amyloid fibrils. We prepared condensates of the lag-series IDPs and initiated monitoring ThT fluorescence immediately after their formation. L20 showed a continuous increase in ThT fluorescence, reaching a plateau around 10 hours, while NP exhibited a ∼10-hour lag phase followed by a gradual increase, reaching a plateau at around 30 hours. L10 maintained a high intensity from the beginning, although no increase was observed in the other lag-series IDPs (Figs. 3A and S5A). We confirmed the presence of amyloid fibrils in the L10, L20, and NP samples by negative staining transmission electron microscopy (TEM) (Fig. 3B), while only aggregates were detected in L30, L40, and L50 (Fig. S5B).

**Fig. 3.**
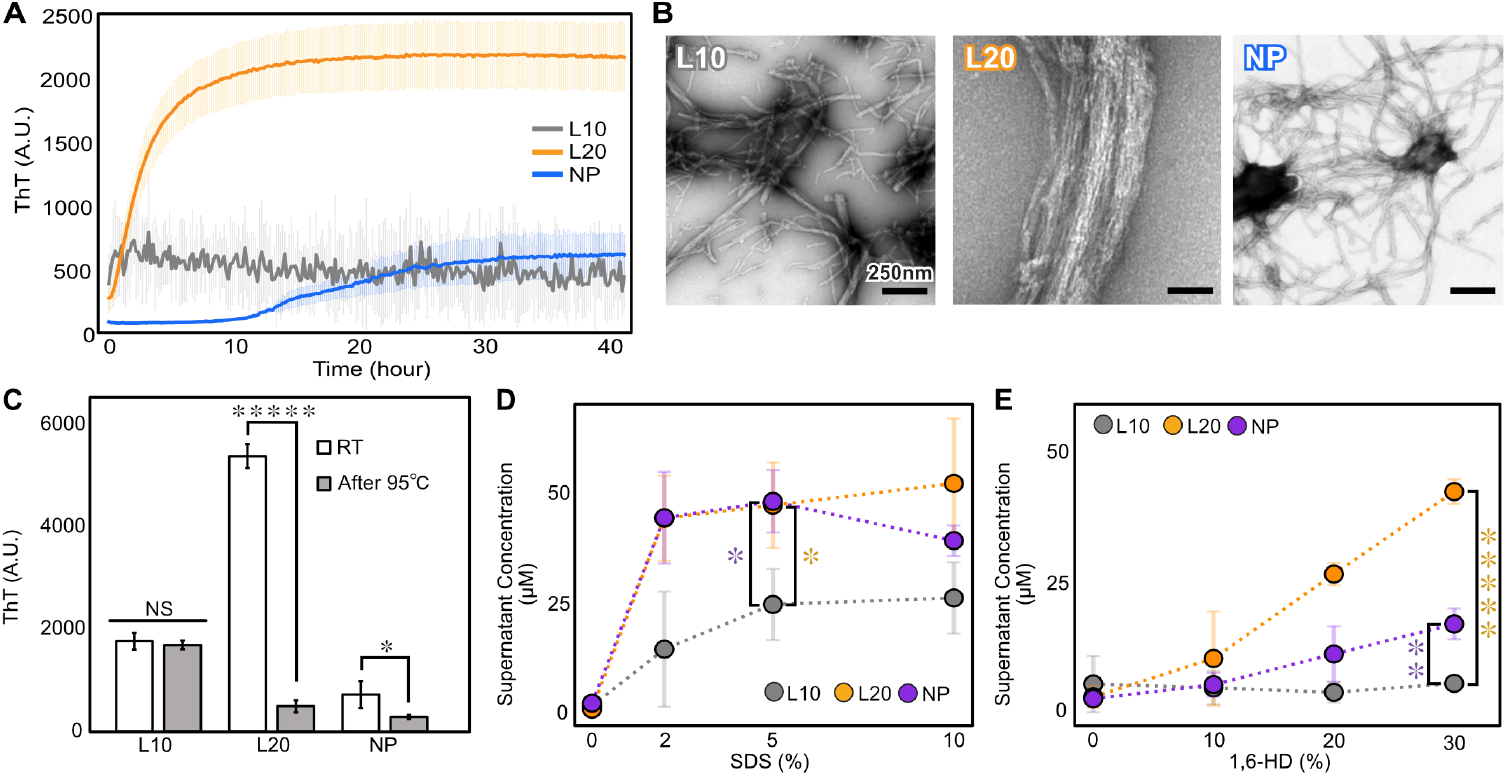
Amyloid fibril formation and condensate stability of lag-series IDPs. **(A)** ThT assay of L10, L20, and NP. Excitation 445 nm, emission 485 nm. Curves represent mean ± SD (N = 3). **(B)** TEM images of L10, L20, and NP. Fibrils were formed by incubating the same solution as that in A at room temperature for 72 hours. Scale bar, 250 nm. **(C)** Heat resistance of L10, L20, and NP condensates. Condensates were prepared as (**B**). Samples were heated at 95 ℃ for 10 min, and ThT fluorescence was measured. Bar plots show mean ± SD (N = 3). RT: unheated; 95 ℃: heated. Asterisks indicate *p*-values by Welch’s t-test: ^*^: *p* < 5.0 × 10^−2, *****^: *p* < 5.0 × 10^−6^, and NS: no significant difference. **(D)** SDS resistance of L10, L20, and NP condensates. Condensates were prepared as (**B**), treated with SDS at indicated concentrations, and incubated 24 hours at room temperature. After centrifugation, protein concentration in supernatants was measured. Data points show mean ± SD (N = 3). For full dataset including WT, see Fig. S5C. One-way ANOVA showed a significant difference at 5 % SDS (*p* = 0.0229). Tukey’s method was used for multiple comparisons. Orange asterisk: L20 vs. L10 (*p* = 0.0376); blue: NP vs. L10 (*p* = 0.0323). **(E)** 1,6-HD resistance of L10, L20, and NP condensates. Same method as (**D**) was used with 1,6-HD. Data points show mean ± SD (N = 3). For the full dataset including WT, see Fig. S5D. One-way ANOVA showed a significant difference at 30 % 1,6-HD (*p* = 2.49 × 10^-6^). Tukey’s method was used for multiple comparisons. Asterisks indicate adjusted *p*-values vs. L10: ^**^: *p* < 5.0 × 10^−3^, and ^*****^: *p* < 5.0 × 10^−6^.

Previous studies have reported that amyloid fibrils matured from condensates can revert to a monomeric state, when exposed to heat or detergent (*44*). To investigate the physical properties of condensates containing amyloid fibrils, we subjected samples of L10, L20, and NP to heat, detergent, and 1,6-HD treatments. Heating at 95 °C for 10 min reduced ThT fluorescence to 9.1 % for L20 and 40 % for NP but had little effect on L10 (Fig. 3C). We also incubated the condensates with sodium dodecyl sulfate (SDS) or 1,6-HD for 24 hours and measured the concentration of solubilized molecules in the supernatant after the centrifugation. In the presence of 5 % SDS, L20 and NP condensates dissolved 94 % and 96 %, respectively, while L10 condensates remained only 52 % dissolved even in 10 % SDS solvent (Fig. 3D). When treated with 1,6-HD, L20 and NP condensates dissolved by 84 % and 33 % at 30 % 1,6-HD, respectively, while L10 condensates showed no dissolution (Fig. 3E). These results suggest that L20 and NP condensates are relatively labile and reversible, whereas L10 condensates exhibit strong resistance to heat and chemical treatment, reflecting its highly stable, possibly irreversible nature.

### Intracellular condensates of lag-series IDPs

Given the distinct fibril-forming propensities of the lag-series IDPs observed in vitro, we next examined whether similar differences would be observed in cells. EYFP-fused L10, L20, NP, and WT in HeLa and COS-7 cells and monitored their localization with a super-resolution fluorescence imaging. The results showed that L20, NP, and WT were diffusively distributed throughout the cytoplasm, whereas L10 accumulated into granule-like foci (Fig. S6). The lack of visible condensates for L20, NP, and WT in cells may reflect the influence of the EYFP or intracellular environment. In contrast, L10 retained its strong tendency to form condensates even under cellular conditions, highlighting its markedly enhanced aggregation property.

### MD simulation analysis of five-residue segments

Lag-series IDPs exhibited distinct condensate properties despite having identical amino acid composition. We hypothesized that these differences arise from sequence-encoded conformational biases in short segments. To evaluate this concept, we performed all-atom molecular dynamics (MD) simulations on five-residue segments, which were generated by sliding a window of five residues along the sequence of each lag-series IDP.

For each segment trajectory, we extracted two parameters (Fig. S7A). (i) detCMD, the determinant of the covariance matrix of the backbone dihedral angles, which represents the conformational variation of the segment. (ii) DMC, the dihedral angle motion correlation, which quantifies the extent of correlation among the backbone dihedral angles of the five residues. A higher DMC indicates cooperative motions of the residues, while a lower DMC indicates independent random motions. We used detCMD as an indicator of conformational variability and DMC as a measure of local dynamic coordination.

As representative examples, L10, L20, and NP are shown in Fig. 4 (for the other IDPs, see Fig. S7). All lag-series IDPs, except for NP, showed periodic patterns in both detCMD and DMC with strong negative correlation (Figs. 4A, S7B and S7E). NP displayed random fluctuations in both parameters but retained a similar negative correlation. These results suggest that five-residue segments with low backbone structure variation move cooperatively due to constrained dihedral angles.

**Fig. 4.**
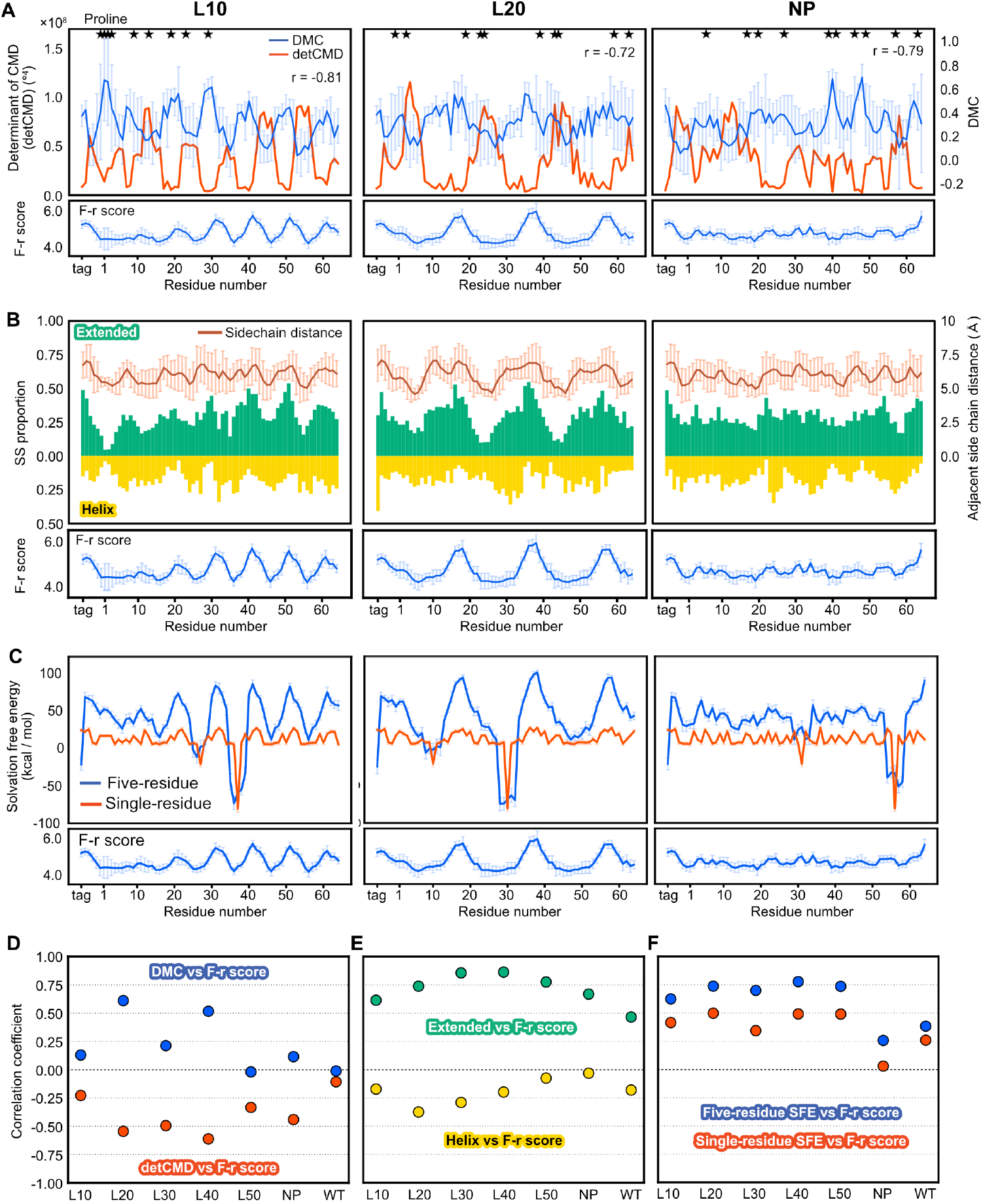
MD simulation analysis of five-residue segments in lag-series IDPs. MD simulations were performed for five-residue segments of lag-series IDPs (see Materials and Methods). Each data point is plotted at the central residue of the corresponding segment (e.g., the segment 1–5 is plotted at residue 3). L30, L40, L50, and WT are shown in Fig. S7. **(A)** Dihedral angle motion correlation (DMC) and determinant of the covariance matrix of the backbone dihedral angles (detCMD). The lower panel shows the corresponding F-r scores. Proline positions are marked by stars. The correlation coefficients between DMC and the detCMD are shown as *r*. **(B)** Secondary structure (SS) proportions and sidechain distances. The lower panel shows the corresponding F-r scores. Green and yellow bars represent extended and α-helical structures, respectively. SS proportions were calculated from the distributions of Ramachandran numbers (Fig. S9). Lines represent the mean values of all sidechain distances of four adjacent residue pairs, with error bars indicating SD. **(C)** Solvation free energy (SFE), with the corresponding F-r scores in the lower panel. Blue lines represent the mean SFE values calculated for five-residue segments, with error bars indicating SD. Red lines represent the mean SFE values calculated for single residues, with error bars indicating SD. **(D)** Correlation coefficients between DMC or detCMD and F-r scores. Blue data points represent correlations between DMC and F-r scores; red data points represent those between detCMD and F-r scores. **(E)** Correlation coefficients between SS proportions and F-r scores. Green data points represent correlations between extended structure proportions and F-r scores; yellow data points represent those between α-helix proportions and F-r scores. **(F)** Correlation coefficients between five-residue or single-residue SFE and F-r scores. Blue data points represent correlations between five-residue SFE and F-r scores; red data points represent those between single-residue SFE and F-r scores.

We next compared detCMD and DMC with the F-r scores. In L20 and L40, DMC correlated positively with the F-r scores, whereas detCMD correlated negatively (Fig. 4D). Similar trends were observed in L10, L30, and NP, but were less apparent for L50 and WT. The weak correlations between the F-r scores and detCMD or DMC may be attributed to proline-rich segments, whose restrained dihedral angles lower detCMD and elevate DMC despite modest F-r scores (Figs. 4A and S7B). Nevertheless, autocorrelation functions of both detCMD and DMC exhibited peaks at the designed lag values (Fig. S8). These results demonstrate that five-residue segments of lag-series IDPs adopt backbone conformational biases as defined by the F-r scores, while also suggest that the F-r scores may capture additional structural features not fully represented by detCMD or DMC.

### Five-residue segments with high F-r scores adopt extended structures including β-strands

Given that detCMD reflects variations in backbone dihedral angles, we asked whether the F-r scores correlate with secondary structure. To test this, we employed the Ramachandran number *R*, a one-dimensional representation of the *ϕ* and *ψ* backbone dihedral angles. *R* values around 0.4 denotes α-helices and *R* values around 0.5 denotes extended structures, including β-strands (*45*). We first analyzed the distribution of *R* values across all segments and observed clear periodic increases and decreases in L10, L20, and L40 (Fig. S9). This observation led us to perform a quantitative analysis, which showed that the proportion of extended structures oscillated with the designed lag values (Figs. 4B and S7C) and correlated strongly with F-r scores (mean r = 0.71), whereas α-helix content correlated only weakly and negatively (Fig. 4E). We further assessed the persistence of extended structures by measuring their longest duration within each trajectory. This duration was highly correlated with the proportion of extended structures (mean r = 0.84, Fig. S10A) and the F-r scores (mean r = 0.74, Fig. S10B). These results suggest that segments with high F-r scores preferentially adopt and retain extended structures, including β-strands.

In extended structures, the sidechains of successive residues point alternately outward in a zig-zag pattern. Because sidechain volume is one of the parameters used for the F-r score calculation, segments with high F-r scores are enriched in bulky residues, such as tryptophan and tyrosine (Figs. 1B and S2). We therefore asked whether bulky neighboring residues might repel one another and keep the segment straight. As a proxy for this effect, we calculated the average distance between the centers of geometry of adjacent residues (Fig. S7A). The average distances correlated positively with the proportions of extended structures (mean r = 0.72, Fig. S7F) and with the F-r scores (mean r = 0.67, Fig. S7F). These results suggest that repulsion between bulky residues in high F-r score segments limits bending and biases the backbone toward persistent, extended conformations.

### Solvation free energy of five-residue segments

To examine whether these segmental features affect intermolecular interactions, we calculated the solvation free energy (SFE) of each segment using 3D-RISM theory. SFE reflects the balance of interactions between the solute and solvent; segments with lower SFE values are more soluble in water and may act as spacers, whereas those with higher SFE values are less soluble and potentially contribute to intermolecular interactions as stickers. While SFE values calculated for single residues (single-residue SFE) showed no clear patterns, SFE values calculated for five residues (five-residue SFE) exhibited periodicity which aligned with the F-r scores (Figs. 4C and S7D). On average, five-residue SFE correlated with the F-r scores at 0.60, compared to 0.36 for single-residue SFE (Fig. 4F). These results suggest that segments with high F-r scores are less soluble, forming hydrophobic patches that facilitate intermolecular interactions.

Taken together, five-residue segments with high F-r scores are likely to adopt extended structures, including β-strands. These conformational tendencies, combined with reduced solubility, support their potential contribution to intermolecular interactions.

### NMR analysis of local dynamics in lag-series IDPs

To complement the MD analysis, we experimentally examined local dynamics and conformational biases of lag-series IDPs using NMR spectroscopy. To examine the local dynamics, we measured longitudinal (R_1_), transverse relaxation rates (R_2_), and heteronuclear NOE (hNOE) of backbone amide groups. These values have been shown to provide valuable insights into local dynamics of IDPs (*46*). Although L10, L30, L40, and L50 did not yield analyzable spectra due to their low solubility, we successfully obtained these parameters for WT, L20, and NP in 12 % 1,6-HD solvent (Fig. S11A).

To confirm that all three IDPs remain monomeric under these conditions, we estimated an apparent rotational correlation time (τ_c_), assuming isotropic tumbling and neglecting internal motions. The apparent τ_c_ values for WT and L20 (3.4 ns) closely matched the theoretical estimate based on the molecular weight (3.7 ns), while NP showed a slightly shorter value (2.5 ns), suggesting a more compact or dynamic conformation but still consistent with a monomeric state.

To assess the relationship between the F-r scores and local dynamics, we compared the relaxation rates with the F-r scores for L20, NP, and WT. L20 exhibited clear oscillations in R_1_, R_2_, and hNOE values with a periodicity of ∼20 residues, while NP showed nearly uniform values (Fig. 5A). Correlation analysis revealed the strongest agreement in L20 (r = 0.59 for R_1_, 0.68 for R_2_, 0.47 for hNOE), followed by WT (r = 0.55, 0.37, 0.50) (Fig. S12A) and NP (r = 0.44, 0.23, 0.39). These results indicate that regions with high F-r scores, especially in L20, tend to exhibit slower backbone dynamics. However, correlations remained modest at the single-residue level, likely due to contributions from long-range interactions or higher-order structures involving segments larger than five residues, which are not captured by the five-residue model.

**Fig. 5.**
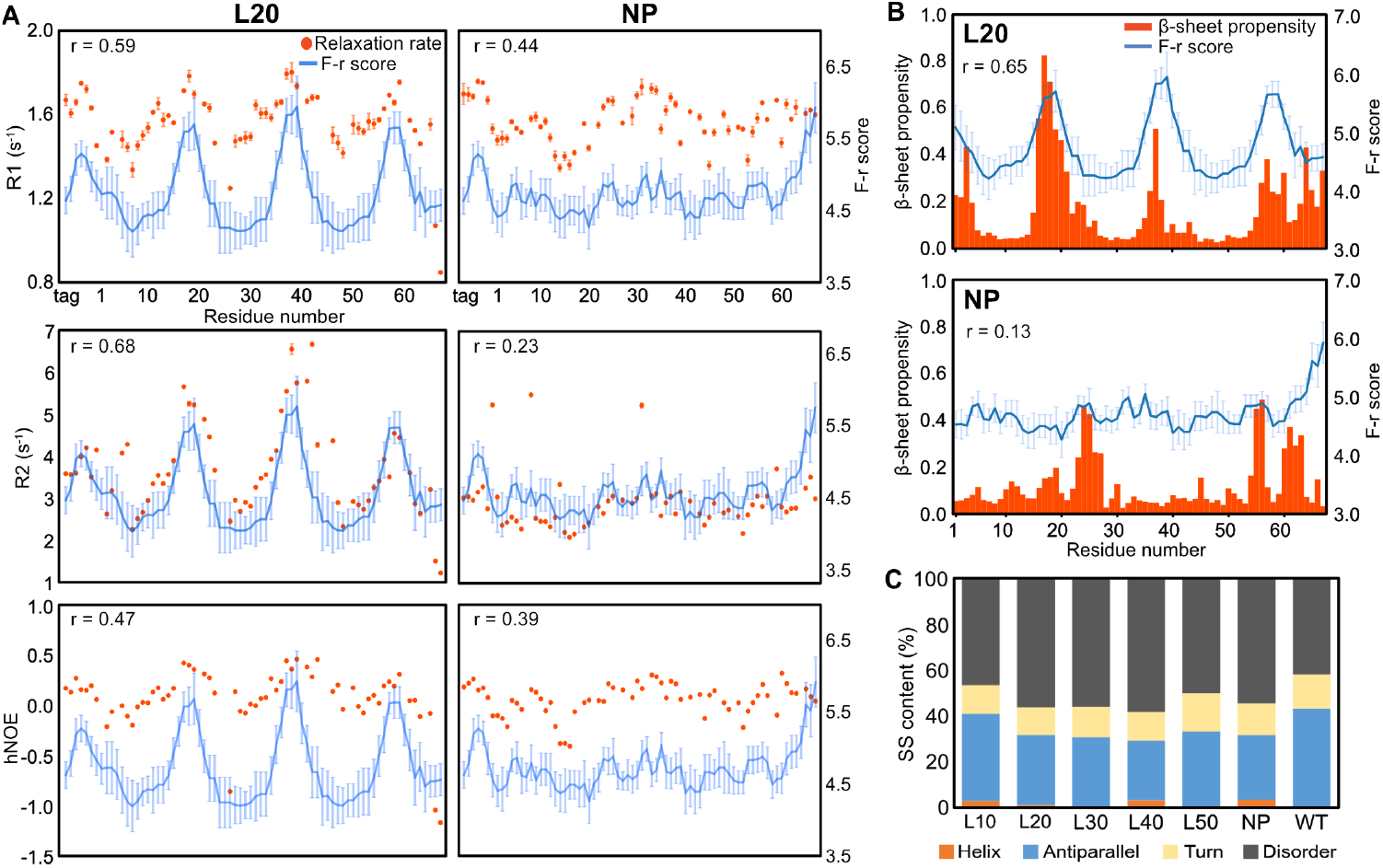
^15^N relaxation rates and secondary structure propensities of L20 and NP. **(A)** R_1_, R_2_ and hNOE of L20 and NP. Relaxation rates were estimated from two time series datasets, and data points represent mean ± SD. The F-r scores are shown in the same way as in Fig. 1B. The correlation coefficients between the relaxation rates and the mean values of F-r scores are shown as *r*. **(B)** β-sheet propensity of L20 and NP. Bar plots represent β-sheet propensity estimated from backbone chemical shifts, and the F-r scores are shown in the same way as in Fig. 1B. The correlation coefficients between the β-sheet propensity and the mean values of F-r scores are shown as *r*. **(C)** Secondary structure (SS) content of lag-series IDPs. SS contents were calculated from CD spectra (Fig. S14). The percentages of each structural element are presented as stacked bar charts.

Compared to our previous study (*38*), the correlation in WT was somewhat weaker. To determine whether this difference was caused by the solvent condition, we repeated the measurements with 10 % 1,6-HD solvent as before (Fig. S11B). However, the relaxation rates remained unchanged (Fig. S12B), suggesting that the discrepancy may arise from differences in instrumentation or experimental setup.

### Lag-series IDPs form β-sheets with distinct structural stabilities

During NMR backbone assignments, we noted that L20 is likely to contain several β-sheet regions. Motivated by this observation, we estimated secondary structure propensities from backbone chemical shifts for L20, NP, and WT. As previously reported (*38*), WT showed uniformly low β-sheet propensity across its sequence (Fig. S12C). NP exhibited dispersed regions with β-sheet propensity over 0.4 (Fig. 5B). In contrast, L20 displayed a clear periodic pattern, with peaks every 20 residues and contiguous stretches exceeding 0.4 in β-sheet propensity, suggesting β-sheet formation in those regions (*47*). None of the three IDPs showed high α-helix propensity (Fig. S13). Notably, the β-sheet-prone regions in L20 corresponded closely with its high F-r score regions (r = 0.65), whereas NP showed no such correlation (r = 0.13).

These findings, together with the MD simulation results, led us to speculate that high F-r score regions may preferentially form β-sheets due to their extended structure bias. Accordingly, as L20, L30, L40, and L50 contain longer high F-r score regions (Fig. S3B), we expect that these IDPs may exhibit higher β-sheet content than NP. To validate this speculation, we conducted circular dichroism (CD) spectroscopy on all lag-series IDPs. Although CD measurements under condensate-forming conditions were hampered by light scattering, they were feasible in 12 % 1,6-HD solution, which is the same condition used for NMR measurements. All lag-series IDPs showed CD spectral features typical of β-sheet structures, characterized by a positive peak at 190– 192 nm and a negative peak at 217–218 nm (Fig. S14) (*48*). To determine secondary structure contents from CD spectra, we used the BeStSel method, which provides accurate secondary structure estimations across diverse protein folds (*49*). Contrary to our expectations that β-sheet structure contents would vary in lag-series IDPs, the results showed that all lag-series IDPs shared similar secondary structure contents, including β-sheet contents (Fig. 5C).

NMR experiments showed a significant difference in β-sheet propensity between L20 and NP, whereas CD experiments indicated that both had nearly identical β-sheet content. We propose that this discrepancy arises from the different aspects of β-structure detected by NMR and CD experiments. NMR detects intramolecular β-sheets that persist on long time scales, typically in the micro-to millisecond range. By contrast, CD provides an ensemble average of backbone structures adopting β-strand conformations and therefore cannot distinguish between transient β-strand states and intramolecular β-sheets. Collectively, the NMR and CD experiments suggest that all lag-series IDPs populate β-strand conformations to a comparable extent, but their stability differs, with long-duration, metastable intramolecular β-sheets present in L20 and NP. Notably, in L20, these regions appear to coincide with high F-r score regions, implying a potential link between conformational biases and metastable intramolecular β-sheet formation.

### F-r score periodicity governs β-sheet formation

Given that amyloid-forming proteins, such as tau and Aβ42/Aβ40, tend to form transient intramolecular β-sheets (*50, 51*), we hypothesized that the metastable intramolecular β-sheets observed in lag-series IDPs may promote the formation of intermolecular β-sheets that assemble into amyloid fibrils. We sought to characterize metastable β-sheet formation in other lag-series IDPs, however their low solubility prevented experimental validation. Instead, we used the structure prediction program AlphaFold2 (AF2) to evaluate their conformational tendencies (*52*). In L20, AF2 predicted several β-sheet regions across the top five ranked models with an average pLDDT (predicted Local Distance Difference Test) score of 36.4 (Fig. 6A). Notably, these predicted β-sheet regions corresponded to the β-sheet regions observed by NMR and high F-r score regions.

**Fig. 6.**
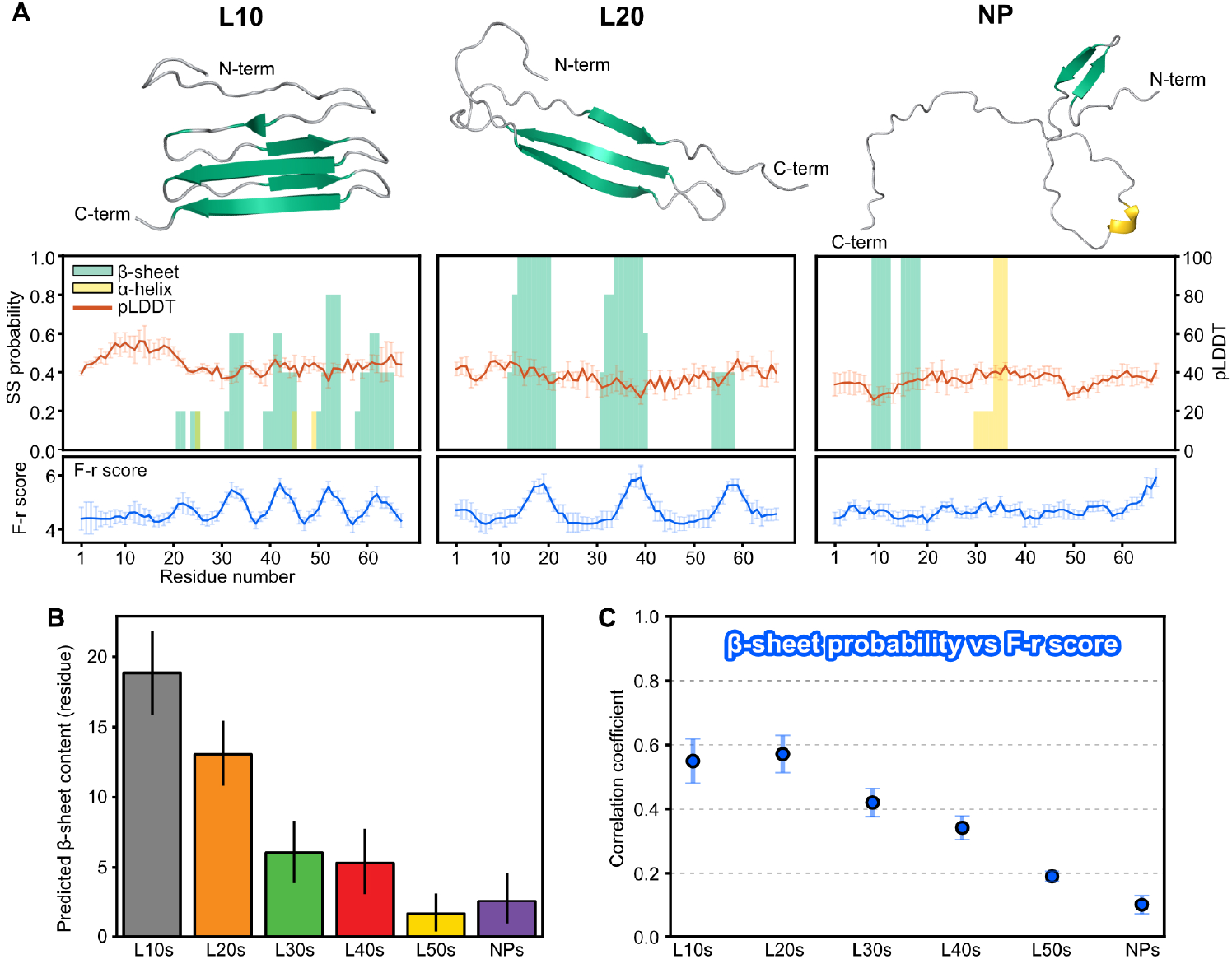
β-sheet propensity of lag-series IDPs including newly designed sequences predicted by AlphaFold2 (AF2) **(A)** AF2 predictions for lag-series IDPs. For each IDP, a representative structure from the top five ranked models is shown at the top. In these structures, β-sheets and α-helices are colored green and yellow, respectively. SS probabilities shown in bar plots were calculated for each residue as the proportion of models exhibiting β-sheet or α-helix conformations among the top five ranked models at each residue. The pLDDT plots represent the mean values across all five models, with error bars indicating SD. The F-r scores are shown in the same way as in Fig. 1B. L30, L40, and L50 are shown in Fig. S15. **(B)** Predicted β-sheet contents for all lag-series IDPs. The predicted β-sheet content was calculated as the expected number of residues forming β-sheet conformations per sequence. Bar plots represent the mean predicted β-sheet contents for each lag-series IDP group, with error bars indicating SD. **(C)** Correlation coefficients between β-sheet probability estimated by AF2 predictions and the F-r scores for each lag-series IDP group. To assess the correspondence between the predicted β-sheet regions and high F-r score regions, the correlation coefficient was calculated between the combined β-sheet probability and the combined F-r scores for 50 different AA-index values. Data points represent the mean value of the correlation coefficients of 50 different F-r scores, with error bars indicating SD.

In contrast, AF2 prediction for NP showed β-sheet regions only in short segments of 9-12 and 15-18 residues with an average pLDDT of 35.5. These regions did not align with β-sheet regions observed by NMR or high F-r score regions. Because pLDDT scores below 50 typically correspond to disordered regions (*53*), these predictions should be interpreted with care. Nonetheless, the agreement between AF2-predicted and NMR-observed β-sheet regions may capture sequence-encoded features of metastable β-sheets within IDPs.

We subsequently applied AF2 to the other lag-series IDPs (Figs. 6A and S15). In L10, the predicted structures revealed periodic β-sheet regions corresponding to high F-r score regions, with an average pLDDT of 42.5. In contrast, L30, L40, and L50 revealed low content of β-sheet regions in dispersed regions, which did not correspond to high F-r score regions. Interestingly, α-helices were predicted for L30, with an average pLDDT of 35.0. In summary, intramolecular β-sheet structures were predicted in high F-r score regions in L10 and L20, while in the other lag-series IDPs, several β-sheet regions were predicted but located outside high F-r score regions. These results suggest that high F-r score regions have intrinsic potential to form β-sheet structures, and that the periodicity of F-r scores, defined by the lag, may play a crucial role in promoting β-sheet formation.

To systematically evaluate the role of the F-r score periodicity in β-sheet formation, we designed 30 new sequences for each lag (180 sequences in total) (Dataset S1, sheet “newIDPsequence”). We performed AF2 predictions for all sequences and quantified their β-sheet content. The results showed that β-sheet content decreased with increased lag (Fig. 6B). To assess the positional alignment between predicted β-sheet regions and high F-r score regions, we calculated correlation coefficients between F-r scores and β-sheet probability. The L10 and L20 groups exhibited strong correlations (r = 0.55 and 0.57) (Fig. 6C), indicating a clear positional preference for β-sheet formation in high F-r score regions. In contrast, these correlations gradually declined as the lags increased in L30, L40, and L50 groups. These results suggest that shorter F-r score periodicity enhances intramolecular β-sheet formation in high F-r score regions.

### Amyloid fibril formation of newly designed lag-series IDPs

To test whether the intramolecular β-sheet tendency revealed by F-r score periodicity translates into amyloid fibril formation, we selected 18 representative sequences from newly designed sequences. For each periodicity group (L10s–L50s), we chose the three variants with the highest ACF(lag^*^) values, and for the NP group we chose the three with the lowest summed ACF(lag) values (Dataset S1, sheet “newIDPsequence”, yellow). Within each group, these sequences are labelled 1st, 2nd, and 3rd in descending ACF(lag^*^) values (ascending summed ACF(lag) value for NP).

In the short-periodicity groups (L10s, L20s, and L30s), at least one sequence in each group such as L10s-1st, L20s-1st, L20s-3rd, and L30s-2nd displayed the typical amyloid fibril features, where the ThT signal increased steadily and fibrils were seen with TEM (Figs. 7A, 7B and 7C). Among them, L10s-1st was the most similar to the original L10 sequence. ThT signal rose immediately to a high and stable level, and the condensate was mostly resistant to 1,6-HD, with only about 13 % becoming soluble (Figs. S16A and S16B). The L20s-1st and L20s-3rd sequences also acted like the original L20 sequence. Their ThT signals increased more slowly, but they still formed fibrils with similar morphologies.

**Fig. 7.**
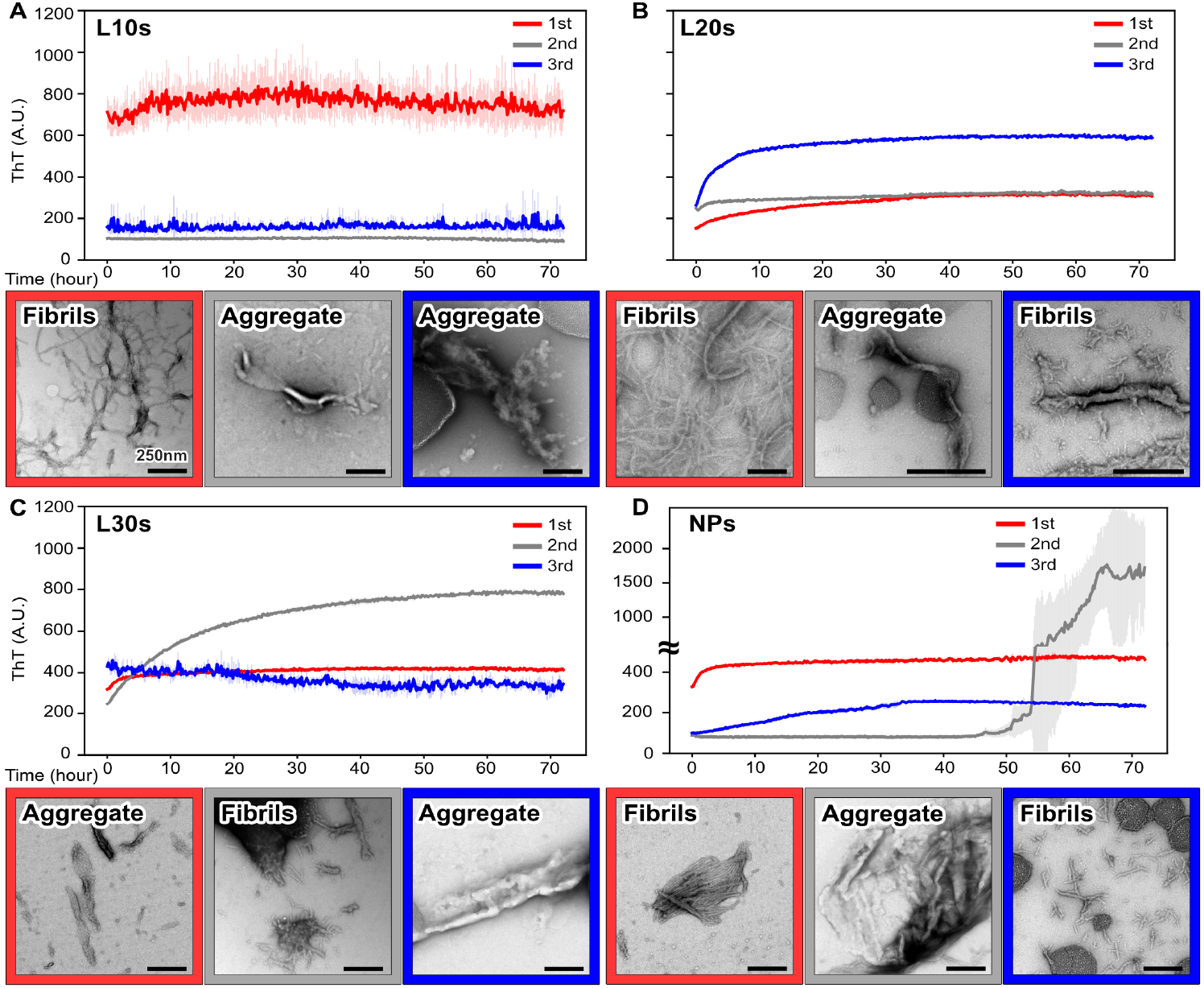
Amyloid fibril formation of newly designed lag-series IDPs. ThT assay and negative staining TEM images for three representative sequences from **(A)** L10s, **(B)** L20s, **(C)** L30s, and **(D)** NPs groups. L40s and L50s are shown in Fig. S17. For ThT assay, excitation 445 nm, emission 485 nm. Curves represent mean ± SD (N = 3). For TEM images, scale bar, 250 nm.

The long-periodicity groups (L40s and L50s) did not show any increase in ThT fluorescence, even though two sequences, L40s-3rd and L50s-2nd, produced fibrils that were visible by TEM (Fig. S17). This pattern, where fibrils are ThT negative but TEM positive, has been reported in time-resolved cryo-EM studies of IAPP-S20G and tau (*54, 55*). These studies observed untwisted fibrils with small, ordered cores during the early stages of assembly, and these immature structures failed to bind ThT but were clearly seen by both TEM and cryo-EM. The fibrils formed by L40s-3rd and L50s-2nd therefore are likely to represent the same immature, unstable form.

The NP group showed diverse behaviors (Fig. 7D). NPs-1st showed only a modest increase in ThT fluorescence, NPs-3rd showed a pronounced increase, and NPs-2nd displayed a sudden increase after about 50 hours yet still yielded no fibrils in TEM. These heterogeneous outcomes imply that, without a defined periodicity, amyloid fibril formation follows several distinct pathways.

Taken together, these findings suggest that a short-periodicity of high F-r score segments (10–30 residues) is a key prerequisite for producing typical amyloid fibrils that bind ThT and are visible by TEM. We propose that this relationship arises from the strong tendency to adopt intramolecular β-sheets, potentially providing a structural basis for subsequent intermolecular β-sheet formation in amyloid fibrils

## Discussion

Our study was motivated by two proposed mechanisms of IDP self-assembly: the sticker-and-spacer framework based on transient point-to-point interactions, and the cross-β hypothesis involving more persistent surface-to-surface interactions. We hypothesized that conformational biases of IDP segments dictate which mode predominates, and that tuning these biases could yield artificial IDPs with distinct behaviors. To test this, we used the F-r model, which quantifies conformational biases in five-residue segments. Guided by this model, we shuffled the TIA-1 PLD sequence while preserving composition, generating lag-series IDPs with distinct periodic patterns. In vitro, all except L10 formed condensates, but with markedly different properties: NP formed large, fluid spherical condensates; L20 showed intermediate behavior; L30, L40, and L50 formed smaller, less soluble condensates; and L10 aggregated irreversibly.

To understand their molecular details, we performed MD simulations on all five-residue segments. These analyses revealed that high F-r score segments tend to exhibit cooperative backbone motion and adopt extended structures, including β-strands. These segments present poorly solvated surfaces, which are conducive to persistent interactions. In contrast, low F-r score segments exhibited uncorrelated, random motions more compatible with transient contacts (Fig. S18). Thus, we surmise that the periodic arrangement of conformationally biased segments may shape the balance between point-to-point and surface-to-surface interactions, which in turn dictates condensate properties. IDPs with more extended high-F-r score regions (e.g., L30, 40, 50) favor dense and persistent associations, while those with few high F-r score regions (e.g., NP) favor dynamic and transient associations. In fact, previous studies have indicated that the segregation of sticker residues, such as aromatic and charged residues, dictates condensate properties (*19, 22, 24*). These results suggest that the more concentrated the stickers, the more dominant the surface-to-surface interactions, whereas the more dispersed the stickers, the more dominant the point-to-point interactions. Thus, we suggest that the F-r model can comprehensively explain the condensate properties by encompassing both modes of interaction.

Among the original lag-series IDPs, L10, L20, and NP demonstrated the capacity to form amyloid fibrils. Further analyses of newly designed lag-series IDPs suggested that lag-series IDP groups with shorter periodicity tended to form typical amyloid fibrils that were ThT-positive and clearly visible by TEM. To probe the molecular basis, we characterized their monomeric states by NMR and CD spectroscopy, complemented with AF2 predictions. We chose this approach because local conformational preferences and intermolecular interactions of IDPs in the condensed phase can be extrapolated from those of monomeric states in the dilute phase (*19, 23– 25*). These structural analyses suggest that the periodicity of F-r scores, that is, the spacing connecting high F-r score regions, is the key determinant of whether metastable intramolecular β-sheets can form. We surmise that high F-r score regions separated by short spacers of roughly 10 to 30 residues can approach one another within a single chain to form metastable β-sheets. Once two molecules encounter one another, these intramolecular β-sheets readily convert into intermolecular pairings that seed amyloid fibril growth. In contrast, longer spacers of 30 residues or more keep high F-r score regions apart and reduce nascent intramolecular β-sheet formation. Homogeneous intermolecular pairing therefore is rare, and any fibril core that does form remains small and labile (Fig. S18). This spacing effect is consistent with intramolecular β-hairpins that accelerate fibril nucleation in tau or Aβ peptides (*50, 51*).

Within the L10 group, sequences such as the original L10 and L10s-1st formed amyloid fibrils rapidly and produced condensates that remained largely insoluble even in 1,6-HD. These properties are consistent with the very short spacers that separate their high F-r score regions. Because these regions are enriched in hydrophobic residues, their close proximity greatly increases the likelihood of direct hydrophobic contacts. Once such contacts form, neighboring regions can stack into a contiguous, water-excluded core, stabilizing the amyloid architecture. Structural studies of pathogenic fibrils support this view; irreversible amyloids typically contain tightly packed hydrophobic cores, whereas reversible fibrils often adopt polar zippers built from glutamine-rich or otherwise polar residues (*56*). By contrast, the longer spacers in L20, L30, and the higher-lag groups insert polar residues between high F-r score regions, reducing hydrophobic burial, favoring polar zipper-like hydrophilic cores and thereby permitting condensate disassembly.

Interestingly, the NP group, including original NP, exhibited amyloid fibril formation with highly heterogeneous behaviors, such as variable lag times and ThT fluorescence intensities. These findings reinforce the idea that NP follows fibril formation mechanisms distinct from those observed in periodic sequences. NP lacks regions acting for surface-to-surface interactions due to randomized F-r scores, and thus its amyloid fibril formation might occur as a result of sampling a wide range of conformations. Among these conformations, it may stochastically adopt structures acting as nuclei for amyloid fibril formation. In the study of Aβ42, it has been suggested that elongation phase of fibril formation needs random search of conformations for reducing time trapped in misregistered conformations (*57–59*). Thus, NP sequences may undergo such a random sampling process compared to the other lag-series IDPs, necessitating further research to fully explain the mechanism underlying amyloid fibril formation.

Our research demonstrates that modifying local conformational biases of IDPs can alter their interaction modes, thereby controlling the condensate properties and fibril formation propensity. We believe that the introduction of the F-r model, which is based on sequence-encoded conformational biases, will have broad applications, including the rational design of IDPs with tuned self-assembly properties and the prediction of phase separation behaviors of uncharacterized IDPs, by bridging the sticker-and-spacer framework and the cross-β hypothesis.

## Materials and Methods

### F-r score calculations

F-r scores were calculated using 50 amino acid indices representing various physicochemical properties. These indices were selected from the AA-index database, which contains 563 parameters (*60*), based on a previous study (*38*). The list of used indices is provided in Dataset S1, sheet “indexvalues”.

Before calculating the F-r scores, each amino acid sequence was converted into a set of 50 corresponding AA-index values. The F-r score was then defined as the geometric mean of AA-index values calculated over a sliding window of five residues. The F-r score for residue *i* and AA-index *j* was calculated using the following formula:

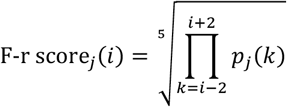

where *p*_*j*_(*k*) represents the AA-index value for residue *k* in AA-index *j*. Each lag-series IDP sequence consisted of 67 residues (excluding tags) and was represented as a 50 × 67 matrix, with rows corresponding to AA-indices and columns to residue numbers. This matrix was shown as a heat map showing the F-r score profile across the sequence (Fig. S3A).

Full experimental details are available in the Supplementary Methods section of the SI appendix.

## Supporting information

Supplementary Methods, Figures, and References

## Acknowledgments

We thank Division of Electron Microscopic Study, Center for Anatomical Studies, Graduate School of Medicine, Kyoto University for technical assistance in electron microscopy. Numerical calculations were partially conducted at the Research Center for Computational Science, Institute for Molecular Science, National Institutes of Natural Sciences (Project: Project 24-IMS-C066 and 25-IMS-C068), the GENKAI supercomputer provided by Research Institute for Information Technology, Kyushu University, MCRP-S at the Center for Computational Science, University of Tsukuba, and supercomputer Fugaku provided by the RIKEN Center for Computational Science (Data-Driven Research Methods Development and Materials Innovation Led by Computational Materials Science: JPMXP1020230327, Project ID: hp230212, hp240223, hp250229). All members of our laboratory, in particular M. Ueno and D. Inaoka, for helpful discussion. This work was supported by MEXT KAKENHI Grant Number JP22H05087 (to N.Y, H.N, N.S.), MEXT KAKENHI Grant Number JP22H05089 (to N.Y.), MEXT KAKENHI Grant Number JP22H05090 (to N.S.), MEXT KAKENHI Grant Number JP22H05091 (to H.N.), JSPS KAKENHI Grant Number JP24K01434 (to N.Y.), MEXT “Data Creation and Utilization-Type Material Research and Development” JPMXP1122714694 (to N.Y.)

## Data and materials availability

All data are available in the main text and the supplementary materials. The code for F-r score calculation and sequence generation is available at https://github.com/ryogakobakoba/Lagseries. Backbone chemical shifts data for NP and L20 have been deposited in Biological Magnetic Resonance Bank (accession numbers: 53269 and 53279). AF2 predicted structure models for all generated sequences, including the original lag-series IDPs, have been deposited in Zenodo (https://doi.org/10.5281/zenodo.15861216).

## References

1. M. E. Oates, P. Romero, T. Ishida, M. Ghalwash, M. J. Mizianty, B. Xue, Z. Dosztányi, N. Uversky, Z. Obradovic, L. Kurgan, A. K. Dunker, J. Gough, D2P2: Database of disordered protein predictions. Nucleic Acids Res. 41, 508–516 (2013).

2. E. Gomes, J. Shorter, The molecular language of membraneless organelles. J. Biol. Chem. 294, 7115–7127 (2019).

3. D. Moses, K. Guadalupe, F. Yu, E. Flores, A. R. Perez, R. McAnelly, N. M. Shamoon, G. Kaur, E. Cuevas-Zepeda, A. D. Merg, E. W. Martin, A. S. Holehouse, S. Sukenik, Structural biases in disordered proteins are prevalent in the cell. Nat. Struct. Mol. Biol., 2021.11.24.469609 (2024).

4. R. K. Das, K. M. Ruff, R. V. Pappu, Relating sequence encoded information to form and function of intrinsically disordered proteins. Curr. Opin. Struct. Biol. 32, 102–112 (2015).

5. K. Sugase, H. J. Dyson, P. E. Wright, Mechanism of coupled folding and binding of an intrinsically disordered protein. Nature 447, 1021–1025 (2007).

6. J. M. Rogers, V. Oleinikovas, S. L. Shammas, C. T. Wong, D. De Sancho, C. M. Baker, J. Clarke, Interplay between partner and ligand facilitates the folding and binding of an intrinsically disordered protein. Proc. Natl. Acad. Sci. U. S. A. 111, 15420–15425 (2014).

7. W. Borcherds, F. X. Theillet, A. Katzer, A. Finzel, K. M. Mishall, A. T. Powell, H. Wu, Manieri, C. Dieterich, P. Selenko, A. Loewer, G. W. Daughdrill, Disorder and residual helicity alter p53-Mdm2 binding affinity and signaling in cells. Nat. Chem. Biol. 10, 1000–1002 (2014).

8. A. S. Krois, H. Jane Dyson, P. E. Wright, Long-range regulation of p53 DNA binding by its intrinsically disordered N-terminal transactivation domain. Proc. Natl. Acad. Sci. U. S. A. 115, E11302–E11310 (2018).

9. S. F. Banani, H. O. Lee, A. A. Hyman, M. K. Rosen, Biomolecular condensates: Organizers of cellular biochemistry. Nat. Rev. Mol. Cell Biol. 18, 285–298 (2017).

10. A. Patel, H. O. Lee, L. Jawerth, S. Maharana, M. Jahnel, M. Y. Hein, S. Stoynov, J. Mahamid, S. Saha, T. M. Franzmann, A. Pozniakovski, I. Poser, N. Maghelli, L. A. Royer, M. Weigert, E. W. Myers, S. Grill, D. Drechsel, A. A. Hyman, S. Alberti, A Liquid-to-Solid Phase Transition of the ALS Protein FUS Accelerated by Disease Mutation. Cell 162, 1066–1077 (2015).

11. S. Ray, N. Singh, R. Kumar, K. Patel, S. Pandey, D. Datta, J. Mahato, R. Panigrahi, A. Navalkar, S. Mehra, L. Gadhe, D. Chatterjee, A. S. Sawner, S. Maiti, S. Bhatia, J. A. Gerez, A. Chowdhury, A. Kumar, R. Padinhateeri, R. Riek, G. Krishnamoorthy, S. K. Maji, α-Synuclein aggregation nucleates through liquid–liquid phase separation. Nat. Chem. 12, 705–716 (2020).

12. Y. Sun, S. Zhang, J. Hu, Y. Tao, W. Xia, J. Gu, Y. Li, Q. Cao, D. Li, C. Liu, Molecular structure of an amyloid fibril formed by FUS low-complexity domain. iScience 25, 103701 (2022).

13. Z. Zhang, G. Huang, Z. Song, A. J. Gatch, F. Ding, Amyloid Aggregation and Liquid– Liquid Phase Separation from the Perspective of Phase Transitions. J. Phys. Chem. B 127, 6241–6250 (2023).

14. J. Brettschneider, K. Del Tredici, V. M. Y. Lee, J. Q. Trojanowski, Spreading of pathology in neurodegenerative diseases: A focus on human studies. Nat. Rev. Neurosci. 16, 109–120 (2015).

15. R. Hervás, L. Li, A. Majumdar, M. del C. Fernández-Ramírez, J. R. Unruh, B. D. Slaughter, A. Galera-Prat, E. Santana, M. Suzuki, Y. Nagai, M. Bruix, S. Casas-Tintó, M. Menéndez, D. V. Laurents, K. Si, M. Carrión-Vázquez, Molecular Basis of Orb2 Amyloidogenesis and Blockade of Memory Consolidation. PLoS Biol. 14, 1–32 (2016).

16. J. Li, T. McQuade, A. B. Siemer, J. Napetschnig, K. Moriwaki, Y. S. Hsiao, E. Damko, D. Moquin, T. Walz, A. McDermott, F. K. M. Chan, H. Wu, The RIP1/RIP3 necrosome forms a functional amyloid signaling complex required for programmed necrosis. Cell 150, 339–350 (2012).

17. C. P. Brangwynne, P. Tompa, R. V. Pappu, Polymer physics of intracellular phase transitions. Nat. Phys. 11, 899–904 (2015).

18. A. C. Murthy, G. L. Dignon, Y. Kan, G. H. Zerze, S. H. Parekh, J. Mittal, N. L. Fawzi, Molecular interactions underlying liquid−liquid phase separation of the FUS low-complexity domain. Nat. Struct. Mol. Biol. 26, 637–648 (2019).

19. E. W. Martin, A. S. Holehouse, I. Peran, M. Farag, J. J. Incicco, A. Bremer, C. R. Grace, A. Soranno, R. V. Pappu, T. Mittag, Valence and patterning of aromatic residues determine the phase behavior of prion-like domains. Science (80-.). 367, 694–699 (2020).

20. J. P. Brady, P. J. Farber, A. Sekhar, Y. H. Lin, R. Huang, A. Bah, T. J. Nott, H. S. Chan, A. J. Baldwin, J. D. Forman-Kay, L. E. Kay, Structural and hydrodynamic properties of an intrinsically disordered region of a germ cell-specific protein on phase separation. Proc. Natl. Acad. Sci. U. S. A. 114, E8194–E8203 (2017).

21. W. Borcherds, A. Bremer, M. B. Borgia, T. Mittag, How do intrinsically disordered protein regions encode a driving force for liquid–liquid phase separation? Curr. Opin. Struct. Biol. 67, 41–50 (2021).

22. A. Bremer, M. Farag, W. M. Borcherds, I. Peran, E. W. Martin, R. V. Pappu, T. Mittag, Deciphering how naturally occurring sequence features impact the phase behaviours of disordered prion-like domains. Nat. Chem. 14, 196–207 (2022).

23. M. Farag, S. R. Cohen, W. M. Borcherds, A. Bremer, T. Mittag, R. V Pappu, Condensates formed by prion-like low-complexity domains have small-world network structures and interfaces defined by expanded conformations. Nat. Commun. 13, 7722 (2022).

24. D. Sundaravadivelu Devarajan, J. Wang, B. Szała-Mendyk, S. Rekhi, A. Nikoubashman, Y. C. Kim, J. Mittal, Sequence-dependent material properties of biomolecular condensates and their relation to dilute phase conformations. Nat. Commun. 15, 1912 (2024).

25. S. Rekhi, C. G. Garcia, M. Barai, A. Rizuan, B. S. Schuster, K. L. Kiick, J. Mittal, Expanding the molecular language of protein liquid–liquid phase separation. Nat. Chem. 16, 1113–1124 (2024).

26. M. Linsenmeier, L. Faltova, C. Morelli, U. Capasso Palmiero, C. Seiffert, A. M. Küffner, D. Pinotsi, J. Zhou, R. Mezzenga, P. Arosio, The interface of condensates of the hnRNPA1 low-complexity domain promotes formation of amyloid fibrils. Nat. Chem. 15, 1340–1349 (2023).

27. M. R. Sawaya, S. Sambashivan, R. Nelson, M. I. Ivanova, S. A. Sievers, M. I. Apostol, M. J. Thompson, M. Balbirnie, J. J. W. Wiltzius, H. T. McFarlane, A. Madsen, C. Riekel, D. Eisenberg, Atomic structures of amyloid cross-β spines reveal varied steric zippers. Nature 447, 453–457 (2007).

28. F. Luo, X. Gui, H. Zhou, J. Gu, Y. Li, X. Liu, M. Zhao, D. Li, X. Li, C. Liu, Atomic structures of FUS LC domain segments reveal bases for reversible amyloid fibril formation. Nat. Struct. Mol. Biol. 25, 341–346 (2018).

29. X. Gui, F. Luo, Y. Li, H. Zhou, Z. Qin, Z. Liu, J. Gu, M. Xie, K. Zhao, B. Dai, W. S. Shin, J. He, L. He, L. Jiang, M. Zhao, B. Sun, X. Li, C. Liu, D. Li, Structural basis for reversible amyloids of hnRNPA1 elucidates their role in stress granule assembly. Nat. Commun. 10 (2019).

30. E. L. Guenther, Q. Cao, H. Trinh, J. Lu, M. R. Sawaya, D. Cascio, D. R. Boyer, J. A. Rodriguez, M. P. Hughes, D. S. Eisenberg, Atomic structures of TDP-43 LCD segments and insights into reversible or pathogenic aggregation. Nat. Struct. Mol. Biol. 25, 463–471 (2018).

31. M. P. Hughes, M. R. Sawaya, D. R. Boyer, L. Goldschmidt, J. A. Rodriguez, D. Cascio, L. Chong, T. Gonen, D. S. Eisenberg, Atomic structures of low-complexity protein segments reveal kinked β sheets that assemble networks. Science (80-.). 359, 698–701 (2018).

32. K. A. Murray, D. Evans, M. P. Hughes, M. R. Sawaya, C. J. Hu, K. N. Houk, D. Eisenberg, Extended β-Strands Contribute to Reversible Amyloid Formation. ACS Nano 16, 2154–2163 (2022).

33. K. A. Murray, M. P. Hughes, C. J. Hu, M. R. Sawaya, L. Salwinski, H. Pan, S. W. French, P. M. Seidler, D. S. Eisenberg, Identifying amyloid-related diseases by mapping mutations in low-complexity protein domains to pathologies. Nat. Struct. Mol. Biol. 29, 529–536 (2022).

34. S. Dey, M. Macainsh, H. X. Zhou, SI: Sequence-Dependent Backbone Dynamics of Intrinsically Disordered Proteins. J. Chem. Theory Comput. 18, 6310–6323 (2022).

35. M. Kato, X. Zhou, S. L. McKnight, How do protein domains of low sequence complexity work? Rna 28, 3–15 (2022).

36. S. L. McKnight, Protein domains of low sequence complexity-dark matter of the proteome. Genes Dev. 38, 205–212 (2024).

37. K. A. Burke, A. M. Janke, C. L. Rhine, N. L. Fawzi, Residue-by-Residue View of In Vitro FUS Granules that Bind the C-Terminal Domain of RNA Polymerase II. Mol. Cell 60, 231–241 (2015).

38. N. Sekiyama, K. Takaba, S. Maki-Yonekura, K. I. Akagi, Y. Ohtani, K. Imamura, T. Terakawa, K. Yamashita, D. Inaoka, K. Yonekura, T. S. Kodama, H. Tochio, ALS mutations in the TIA-1 prion-like domain trigger highly condensed pathogenic structures. Proc. Natl. Acad. Sci. U. S. A. 119, 1–12 (2022).

39. N. Sekiyama, R. Kobayashi, T. S. Kodama, Toward a high-resolution mechanism of intrinsically disordered protein self-assembly. J. Biochem. 174, 391–398 (2023).

40. A. Abyzov, N. Salvi, R. Schneider, D. Maurin, R. W. H. Ruigrok, M. R. Jensen, M. Blackledge, Identification of Dynamic Modes in an Intrinsically Disordered Protein Using Temperature-Dependent NMR Relaxation. J. Am. Chem. Soc. 138, 6240–6251 (2016).

41. N. Salvi, A. Abyzov, M. Blackledge, Multi-Timescale Dynamics in Intrinsically Disordered Proteins from NMR Relaxation and Molecular Simulation. J. Phys. Chem. Lett. 7, 2483–2489 (2016).

42. N. Salvi, A. Abyzov, M. Blackledge, Analytical Description of NMR Relaxation Highlights Correlated Dynamics in Intrinsically Disordered Proteins. Angew. Chemie - Int. Ed. 56, 14020–14024 (2017).

43. S. Guseva, V. Schnapka, W. Adamski, D. Maurin, R. W. H. Ruigrok, N. Salvi, M. Blackledge, Liquid–Liquid Phase Separation Modifies the Dynamic Properties of Intrinsically Disordered Proteins. J. Am. Chem. Soc. 145, 10548–10563 (2023).

44. I. Peran, T. Mittag, Molecular structure in biomolecular condensates. Curr. Opin. Struct. Biol. 60, 17–26 (2020).

45. R. V. Mannige, J. Kundu, S. Whitelam, The Ramachandran Number: An Order Parameter for Protein Geometry. PLoS One 11, 1–14 (2016).

46. A. C. Murthy, N. L. Fawzi, The (un)structural biology of biomolecular liquid-liquid phase separation using NMR spectroscopy. J. Biol. Chem. 295, 2375–2384 (2020).

47. Y. Shen, A. Bax, Protein backbone and sidechain torsion angles predicted from NMR chemical shifts using artificial neural networks. J. Biomol. NMR 56, 227–241 (2013).

48. N. Greenfield, G. D. Fasman, Computed Circular Dichroism Spectra for the Evaluation of Protein Conformation. Biochemistry 8, 4108–4116 (1969).

49. A. Micsonai, F. Wien, É. Bulyáki, J. Kun, É. Moussong, Y. H. Lee, Y. Goto, M. Réfrégiers, J. Kardos, BeStSel: A web server for accurate protein secondary structure prediction and fold recognition from the circular dichroism spectra. Nucleic Acids Res. 46, W315–W322 (2018).

50. L. Sari, S. Bali, L. A. Joachimiak, M. M. Lin, Hairpin trimer transition state of amyloid fibril. Nat. Commun. 15 (2024).

51. S. G. Itoh, M. Yagi-Utsumi, K. Kato, H. Okumura, Key Residue for Aggregation of Amyloid-β Peptides. ACS Chem. Neurosci. 13, 3139–3151 (2022).

52. J. Jumper, R. Evans, A. Pritzel, T. Green, M. Figurnov, O. Ronneberger, K. Tunyasuvunakool, R. Bates, A. Žídek, A. Potapenko, A. Bridgland, C. Meyer, S. A. A. Kohl, A. J. Ballard, A. Cowie, B. Romera-Paredes, S. Nikolov, R. Jain, J. Adler, T. Back, S. Petersen, D. Reiman, E. Clancy, M. Zielinski, M. Steinegger, M. Pacholska, T. Berghammer, S. Bodenstein, D. Silver, O. Vinyals, A. W. Senior, K. Kavukcuoglu, P. Kohli, D. Hassabis, Highly accurate protein structure prediction with AlphaFold. Nature 596, 583–589 (2021).

53. J. L. Binder, J. Berendzen, A. O. Stevens, Y. He, J. Wang, N. V. Dokholyan, T. I. Oprea, AlphaFold illuminates half of the dark human proteins. Curr. Opin. Struct. Biol. 74, 102372 (2022).

54. S. Lövestam, D. Li, J. L. Wagstaff, A. Kotecha, D. Kimanius, S. H. McLaughlin, A. G. Murzin, S. M. V. Freund, M. Goedert, S. H. W. Scheres, Disease-specific tau filaments assemble via polymorphic intermediates. Nature 625, 119–125 (2024).

55. M. Wilkinson, Y. Xu, D. Thacker, A. I. P. Taylor, D. G. Fisher, R. U. Gallardo, S. E. Radford, N. A. Ranson, Structural evolution of fibril polymorphs during amyloid assembly. Cell 186, 5798-5811.e26 (2023).

56. M. R. Sawaya, M. P. Hughes, J. A. Rodriguez, R. Riek, D. S. Eisenberg, The expanding amyloid family: Structure, stability, function, and pathogenesis. Cell 184, 4857–4873 (2021).

57. Z. Jia, J. D. Schmit, J. Chen, Amyloid assembly is dominated by misregistered kinetic traps on an unbiased energy landscape. Proc. Natl. Acad. Sci. U. S. A. 117, 10322– 10328 (2020).

58. C. Leonard, C. Phillips, J. McCarty, Insight Into Seeded Tau Fibril Growth From Molecular Dynamics Simulation of the Alzheimer’s Disease Protofibril Core. Front. Mol. Biosci. 8, 1–13 (2021).

59. C. Bleiholder, N. F. Dupuis, T. Wyttenbach, M. T. Bowers, Ion mobility–mass spectrometry reveals a conformational conversion from random assembly to β-sheet in amyloid fibril formation. Nat. Chem. 3, 172–177 (2011).

60. S. Kawashima, AAindex: Amino Acid index database. Nucleic Acids Res. 28, 374–374 (2000).

