## Supplementary Methods, Figures, and References for "Sequence-encoded conformational biases shape self-assembly modes of intrinsically disordered proteins"

**This PDF file includes:**

Supplementary Methods
Supplementary Figures 1 to 18
SI References

**Other Supplementary Materials for this manuscript include the following:**

Supplementary Data
Dataset S1

### Supplementary Methods

#### Designing lag-series IDP sequences

To design amino acid sequences with periodic F-r scores, the WT sequence was given as the starting sequence for this procedure. First, the WT sequence was randomly shuffled to generate 30 new sequences (denoted as WT shuffled sequences). This step provided a diverse set of randomized sequences as a starting point for further analysis. The shuffled sequences were converted into 50 different AA-index values, and the F-r scores were then calculated using the formula described above for each sequence.

Next, to validate periodic patterns of the F-r scores, the autocorrelation functions $\mathrm{ACF}(lag)$ of the F-r scores were calculated, where lag represents the periodicity of the F-r score for lag residues. For the ACF(lag) calculation, a Python module from the statsmodels package version 1.12.0 was used (*1*). The ACF(lag) for AA-index $j$ is calculated using the following formula:

$$\mathrm{ACF}_j(\mathrm{lag}) = \frac{\sum_{i=1}^{\mathrm{N-lag}}\left(\text{F-r score}_j(i) - \mu_j\right)\left(\text{F-r score}_j(i+\mathrm{lag}) - \mu_j\right)}{\sum_{i=1}^{N}\left(\text{F-r score}_j(i) - \mu_j\right)^2}$$

where $\text{F-r score}_j(i)$ represents the F-r score for residue $i$ and AA-index $j$, $\mu_j$ represents the mean value of all F-r scores for AA-index $j$, and $N$ represents the total number of residues, which is 67 for lag-series IDPs.

To extract sequences with higher periodicity of the F-r scores, the $\mathrm{ACF}_j(\mathrm{lag})$ values were averaged across all AA-indices using the following formula:

$$\mathrm{ACF}_{\mathrm{ave}}(\mathrm{lag}^*) = \frac{1}{P}\sum_{\mathrm{j\in AA\text{-}indices}}\left(\mathrm{ACF}_j(\mathrm{lag}^*)\right)$$

Here, $P$ is the total number of AA-indices (= 50), and the ACF values for all AA-indices at a specific lag value $lag^*$ are averaged. The top five sequences with the highest $\mathrm{ACF}_{\mathrm{ave}}(\mathrm{lag}^*)$ values were selected from the WT shuffled sequences. In this study, the $lag^*$ values were set to 10, 20, 30, 40, and 50 residues, respectively.

For the selected sequences, two randomly selected residues were permutated to generate 30 new sequences. This process was applied to each of the five selected sequences, resulting in a total of 150 new sequences (Shuffled sequences). These 150 sequences served as the starting sequences for the next iteration of the F-r score calculations and two residues permutation. This process continued until the $\mathrm{ACF}_{\mathrm{ave}}(\mathrm{lag}^*)$ value reached a plateau. At the final iteration, the top five sequences with the highest $\mathrm{ACF}_{\mathrm{ave}}(\mathrm{lag}^*)$ value were obtained, and one sequence with the highest $\mathrm{ACF}_{\mathrm{ave}}(\mathrm{lag}^*)$ among five sequences was selected as the completed sequence with the strongest periodicity at $\mathrm{lag}^*$.

To generate non-periodic sequence, the $\mathrm{ACF}_j(\mathrm{lag})$ values were summed across all AA-indices and all residues using the following formula:

$$\mathrm{ACF_{sum}} = \sum_{j \in \text{AA-indices}} \sum_{i=1}^{N} \left(\mathrm{ACF}_j(\mathrm{lag})\right)$$

where $i$ represents the residue number, $j$ represents the AA-index, and $N$ represents the total number of residues, which is 67 for lag-series IDPs. The basic procedure for sequence selection is similar to that of the other lag-series IDPs, but in this case, the bottom five sequences with the lowest $\mathrm{ACF_{sum}}$ value were selected from the shuffled sequences. This process was iterated in the same manner as for other sequences, resulting in the final sequence with the weakest periodicity of F-r scores across the entire sequence.

**Plasmid constructs of lag-series IDPs**

DNA fragments coding lag-series IDP sequences were synthesized by GENEWIZ (South Plainfield, NJ, USA), and each fragment was inserted into pET28 vector using the In-Fusion HD Cloning Kit (Takara Bio, Shiga, Japan). All constructs contained an N-terminal $His_{12}$-tag followed by a TEV (Tobacco Etch Virus) protease cleavage site (sequence: ENLYFQG). The sequences of Lag-series IDPs are shown below.

WT: His-TEV-TIA1 PLD
MGSSHHHHHHHHHHHHSENLYFQGGQYVPNGWQVPAYGVYGQPWSQQGFNQ
TQSSAPWMGPNYSVPPPQGQNGSMLPSQPAGYRVAGYETQ

L10: His-TEV-Lag10
MGSSHHHHHHHHHHHHSENLYFQGPPPPGTQTQPAGLPGSNQQPAMFPGSNRQ
PAYWYGSNEQVAYWYGGNSQVQVWYGGSSQVQVYMGQP

L20: His-TEV-Lag20
MGSSHHHHHHHHHHHHSENLYFQGPMVPGGSASQRNQAQVYYFPYSGPPGSG
NQENQTQVWYWPYGAPPGSGNQQSQTQVLYWPVGAPMGQ

L30: His-TEV-Lag30
MGSSHHHHHHHHHHHHSENLYFQGNMPSGGSSQTPPQPARFYQWLGVVYQPY
AQNGESGGSSQTPPQPAQWYQWVGVVYQPYAQNGNPGGM

L40: His-TEV-Lag40
MGSSHHHHHHHHHHHHSENLYFQGQQGNGPQGGGSQQSPPNAYPWYYWVVA
VQPNLSMQPTMTRAQGEGPQGGGSQQSPPNAYPWYYFVVS

L50: His-TEV-Lag50
MGSSHHHHHHHHHHHHSENLYFQGQPEGGGGVYYWFLQSSSNPVQAQPQVPTP
QMNAPQMSYGPTRAQPGYAQPQPNGGGGVYYWWVQSSN

NP: His-TEV-NP
MGSSHHHHHHHHHHHHSENLYFQGGNMAQLPGQVQMGGQSVPGSPSYQVNQP
SYGRYASNWGQPQPAGTYPQQPVGNQFVEPGTYSYPAWW

For mammalian cell experiments, the coding regions of L10, L20, NP and WT were subcloned into the pcDNA3.1 vector between BamHI and XhoI sites using the In-Fusion HD Cloning Kit.

**Protein expression and purification**

*Escherichia coli* BL21 (DE3) was transformed with the appropriate plasmids and cultured in Luria-Bertani (LB) or M9 minimal medium at 37 °C overnight with shaking at 200 rpm. For isotope-labeled samples used in NMR experiments, M9 medium was supplemented with 1 g/L $^{15}NH_4Cl$ and 2 g/L $^{13}$C-D-glucose. Protein expression was induced at an $OD_{600}$ of 1.5 by the addition of 0.1 mM isopropyl-β-D-thiogalactopyranoside (IPTG), followed by incubation at 18 °C for 24 hours. Cells were harvested by centrifugation and washed with PBS.

Cell pellets were resuspended in lysis buffer (50 mM Tris-HCl pH 7.5, 300 mM NaCl and 5 mM 2-mercaptetahnol) supplemented with 0.2 % Triton-X, and 1 mM phenylmethylsulfonyl fluoride (PMSF). Cells were lysed by sonication on ice using an ultrasonic processor (Qsonica). Lysates were centrifuged at 15,000 rpm for 40 min at 4 °C to collect insoluble fractions containing the lag-series IDPs. After discarding the supernatants, the insoluble fractions were solubilized in denaturing buffer (50 mM Tris-HCl pH 8.0 and 8 M Urea) using a homogenizer.

The solubilized samples were centrifuged again at 15,000 rpm for 40 min at 4 °C to remove residual insoluble material. The supernatants were loaded on to a Ni-NTA agarose column (FUJIFILM Wako, Osaka, Japan) equilibrated with denaturing buffer and incubated at room temperature for 2 hours. After discarding the flow-through, the column was washed with 5 column volumes (CV) of denaturing buffer, followed by 15 CV of wash buffer (50 mM Tris-HCl pH 8.0, 8 M Urea and 5 mM Imidazole). Proteins were eluted with 8 CV of elution buffer (50 mM Tris-HCl pH 8.0, 8 M Urea and 250 mM Imidazole).

The eluates were concentrated to 3mL using an Amicon Ultra centrifugal filter (Merck, Darmstadt, Germany) with a 3,000 molecular weight cutoff. To remove urea, the concentrated solutions were dialyzed twice against 0.1 % Trifluoroacetic acid (TFA) overnight at room temperature. The dialyzed solutions were then lyophilized and dissolved in dimethyl sulfoxide (DMSO) or 0.1 % TFA. Samples for NMR were dissolved in 0.1 % TFA; all other samples were dissolved in DMSO. Each sample was adjusted to a final concentration of 3 mM and stored at –80 °C.

**Condensate sample preparation**

Protein stocks were thawed at 95 °C for 1 hour to ensure complete conversion to the monomeric state, then cooled to room temperature. The stocks were subsequently diluted to the desired concentrations in PS (phase separation) buffer (50 mM MES pH 6.5, 150 mM NaCl and 1 mM dithiothreitol). Prior to each experiment, the diluted samples were heated at 95 °C for 10 min and then cooled to room temperature for 30 min. This procedure was based on a previously established protocol (*2*). The initial heating step of the stock samples was critical for experimental reproducibility. The second heating step was implemented to dissolve pre-existing condensates and reset the system. To assess whether heat treatment induced aggregation, we compared condensate ratios of WT samples prepared with and without the heating steps. No significant differences were observed, indicating that the heat treatments did not promote aggregation.

**Microscopy imaging of lag-series IDP condensates**

Condensate samples were prepared at a protein concentration of 25 μM in PS buffer at room temperature, following the condensate sample preparation procedure. For microscopy analysis, a 100 μL condensate sample was placed on a glass-bottom dish (Matsunami Glass Ind., Ltd., Osaka, Japan) and incubated for 2 min to allow the condensates to settle at the bottom.

Bright-filed microscopy was performed using an IX71 microscope (Olympus Corporation, Tokyo, Japan) equipped with a UPLXAPO100XOPH 100x/1.45 oil immersion objective lens (Olympus) and an ORCA-Flash 4.0 CCD camera (Hamamatsu Photonics K.K., Hamamatsu, Japan). Images were acquired using HC-image software (Hamamatsu Photonics).

For fluorescence microscopy, condensates were visualized by adding 5 μM Thioflavin T (ThT) prior to imaging. ThT fluorescence was detected using a NIBA filter set (excitation: 470 - 490 nm, emission: 515 - 550 nm) with the same microscope set up as used for bright-field imaging.

Quantitative analysis of condensate sizes was performed was performed using fluorescence images and the particle analysis module in Fiji (*3*). At least 9 images from different fields were analyzed. Condensates were detected using a minimum area threshold of 0.3 μm², and fused condensates were separated by watershed segmentation. This analysis was applied to the images of L20, L40, L50, NP, and WT.

**Condensate ratio measurement**

Condensate samples were prepared at a protein concentration of 50 μM in PS buffer containing final concentrations of 0 %, 3 %, 6 %, 9 %, 12 %, and 15 % 1,6-hexanediol (1,6-HD), following the condensate sample preparation procedure. A 300 μL aliquot of each sample was placed in a black 96-well plate (Greiner Bio-One, Kremsmünster, Austria) and the tryptophan fluorescence intensity, $F_{drop}$, was measured using a SpectraMaxM2 RF Microplate reader (Molecular Devices, LLC, San Jose, CA, USA) with excitation at 280 nm and an emission at 360 nm. The $F_{drop}$ value represents the total protein content in the solution.

After the initial measurement, samples were transferred to microcentrifuge tubes and centrifuged at 20,400 g for 30 min at room temperature to pellet the condensates. From each tube, 300 μL of supernatant was carefully collected to avoid disturbing the pellet and transferred to a new plate. Fluorescence intensity of the supernatant, $F_{sup}$, was measured using the same settings. The $F_{sup}$ value represents the amount of protein remaining soluble. The condensate ratio, representing the fraction of protein incorporated into condensates, was calculated using the following formula:

$$\text{Condensate ratio} = \frac{F_{con} - F_{sup}}{F_{con}}$$

All measurements were performed in triplicate, and the results were presented as mean ± standard deviation.

**$C_{sat}$ measurement**

To determine the saturation concentration ($C_{sat}$), condensate samples were prepared at a protein concentration of 10 µM in PS buffer, following the condensate sample preparation procedure. Initial attempt without 1,6-HD resulted in protein concentrations in the supernatant falling below the detection limit, preventing accurate $C_{sat}$ determination. Therefore, 8 % 1,6-HD was added to partially dissolve the condensates, allowing quantification of protein in the supernatant. A 150 µL aliquot of each sample was centrifuged at 20,400 g for 30 min to pellet the condensates. From each tube, 150 µL of supernatant was carefully collected without disturbing the pellet. The supernatant protein concentration, representing the $C_{sat}$ value, was measured by absorbance at 280 nm using a NanoPhotometer P-class (Implen, Munich, Germany). Protein concentrations were calculated using the theoretical extinction coefficients 26,930 $M^{-1}$ $cm^{-1}$ estimated from the amino acid sequence. All measurements were performed in triplicate, and the results were presented as mean ± standard deviation.

**ThT assay**

Condensate samples were prepared at a protein concentration of 25 µM in PS buffer, following the condensate sample preparation procedure. A 150 µL aliquot of each sample, containing 5 µM Thioflavin T (ThT), was transferred to a black 96-well plate (Greiner Bio-One). ThT fluorescence was monitored over 40 hours at 30˚C using a SpectraMaxM2 RF Microplate reader (Molecular Devices) with excitation at 445 nm and emission at 485 nm. Fluorescence readings were taken every 10 min, with 5 seconds of shaking prior to each measurement. Fluorescence values from buffer samples containing 1.25 µl DMSO were subtracted from each measurement to account for background signal. The volume of DMSO added was equal to that of the stock sample used to prepare the condensate samples. All measurements were performed in triplicate, and the results were presented as mean ± standard deviation.

**Negative staining TEM**

Condensate samples were prepared at a protein concentration of 25 µM in PS buffer, following the standard condensate preparation protocol. Samples were incubated at room temperature for 72 hours to allow amyloid fibril formation. After incubation, 5 µL of each sample was applied to an ELS-C10 grid (Ohken Shoji, Ltd., Tokyo, Japan) and incubated for 1 minute. The excess solution was removed using filter paper, and the grid was stained with 3 µL of 1% uranyl acetate. After 1 minute, excess stain was removed, and the grid was air-dried overnight. The prepared grids were examined using a JEM-1400 Flash transmission electron microscope (JEOL, Ltd., Tokyo, Japan) operated at an acceleration voltage of 80 kV.

**Thermal stability assay of condensates**

Condensate samples were prepared at a protein concentration of 50 µM in PS buffer containing 5 µM ThT, following the condensate sample preparation procedure. Samples were incubated at room temperature for 72 hours to allow amyloid fibril formation. For each protein, two samples were prepared. After incubation, one sample was heated at 95 °C for 10 min and then cooled to room temperature for 30 min, while the other sample was left untreated. A 300 µL aliquot of each sample was placed in a black 96-well plate

(Greiner Bio-One). ThT fluorescence was measured using a SpectraMaxM2 RF Microplate reader (Molecular Devices) under the same conditions as described in the ThT assay. All measurements were performed in triplicate, and the results were presented as mean ± standard deviation.

**Chemical stability assay of condensates**

Condensate samples were prepared at a protein concentration of 50 μM in PS buffer, following the condensate sample preparation procedure. Samples were incubated at room temperature for 72 hours to allow amyloid fibril formation.

For the stability assay, 60 μL of each sample was centrifuged at 20,400 g for 30 min to pellet the amyloid fibrils. After carefully removing the supernatant, the pellets were resuspended in 60 μL of PS buffer containing 0 %, 10 %, 20 % and 30 % 1,6-HD for the 1,6-HD resistance assay, or 60 μL water containing 0 %, 2 %, 5 % and 10 % of sodium dodecyl sulfate (SDS) for the SDS resistance assay. Samples were mixed well and incubated for 24 hours at room temperature.

After incubation, samples were centrifuged again at 20,400 g for 30 min to sediment any remaining fibrils. A 60 μL aliquot of the supernatant was carefully collected, and protein concentration was determined by absorbance at 280 nm using a NanoPhotometer P-class (Implen), following the same method described in the $C_{sat}$ assay. All measurements were performed in triplicate, and the results were presented as mean ± standard deviation.

**Transient expression of lag-series IDPs in cells**

HeLa cells and COS-7 cells (both from RIKEN BRD, Saitama, Japan) were cultured in high-glucose DMEM (Wako, Osaka, Japan) supplemented with 10% FBS (Sigma) and 1% v/v penicillin-streptomycin (Wako, Osaka, Japan) at 37 °C in a 5% $CO_2$ incubator. DNA sequences for the four lag-series IDPs (L10, L20, NP, and WT) lacking His-tag and TEV cleavage site were amplified by PCR to subclone the sequences into the pEYFP-C1 vector between XhoI and EcoRI sites, in-frame with EYFP. The resultant plasmids encoding N-terminally EYFP-tagged IDPs were transfected to the cell line of interest using Lipofectamine 2000 (Thermo Fisher Scientific, Waltham, MA, USA), according to the manufacturer's instructions. Cells were imaged 24 hours post-transfection with a Leica sp8 confocal fluorescence microscope supplemented with a white excitation laser, using the pre-set EYFP optical setup and a deconvolution-based super-resolution mode, Lightning.

**MD simulation**

To investigate conformational biases of lag-series IDPs at the segment level, all-atom MD simulations were performed on five-residue segments. These segments were generated by sliding a window of five residues along the sequence of each lag-series IDP. The initial segment was $^{18}$ENLYF$^{22}$ (omitting the His tag), and a total of 70 segments were generated across all lag-series IDPs. The N- and C-termini of each segment were capped with acetyl (ACE) and N-methyl (NME) amide groups, respectively.

MD simulations were conducted using Amber20 (*4*). Amber FF99SB force field and OPC parameters were used for the peptide segments and water, respectively (*5*, *6*). Charged amino acids were neutralized with NaCl. After energy minimization, each system underwent 1 ns equilibration followed by 100 ns production run at 300K, 1bar in the NPT ensemble under the Langevin thermostat and Monte Carlo barostat. The time step was 2 fs, and the hydrogen bond length was fixed using the SHAKE method (*7*). The box size was about $50^3$ Å$^3$. A total of 100 snapshots were extracted at 1ns intervals from the 100ns production run for further analysis.

**MD data analysis: DMC and detCMD**

MD trajectories were analyzed using MDAnalysis (*8–10*). The dihedral angles $\phi$ and $\psi$ of each segment were obtained from 100 snapshot structures in the trajectories. To analyze the conformational biases and local dynamics of each segment, two quantities were calculated for five-residue segments: the dihedral angle motion correlation (DMC) and the determinant of the covariance matrix of the backbone dihedral angles (detCMD).

DMC quantifies the extent of correlation among the backbone dihedral angles of the five residues throughout the trajectory. DMC values were calculated as correlation coefficients between the combined $\phi$ and $\psi$ angles of two residues across all snapshots. For each residue, a motion time series (MTS) was constructed using sine and cosine components of the $\phi$ and $\psi$ angles from 100 snapshots:

$$MTS(res) = (v_1, v_2, \cdots v_{100})$$

where $v_i$ are the sine and cosine components of the $\phi$ and $\psi$ angles in snapshot $i$, defined as:

$$v_i = (\sin\phi_i, \cos\phi_i, \sin\psi_i, \cos\psi_i).$$

DMC values were calculated as the Pearson correlation coefficient between the motion time series of all possible residue pairs within each segment (10 pairs in total: 1-2, 1-3, 1-4, 1-5, 2-3, 2-4, 2-5, 3-4, 3-5, 4-5):

$$DMC = Corr[MTS(res\_i), MTS(res\_j)]$$

where $Corr$ denotes the Pearson correlation coefficient, and $res\_i \neq res\_j$.

detCMD provides a single numerical value estimating the overall magnitude of variation in the backbone dihedral angles within a five-residue segment, representing its conformational variability. For each segment, detCMD values were calculated from 500 dihedral angle pairs. Specifically, 100 $\phi$ and 100 $\psi$ angles were extracted from 100 snapshots for each residue, resulting in 500 pairs of dihedral angles (5 residues × 100 snapshots), corresponding to 500 points on the Ramachandran plot.

To quantify the two-dimensional variance in the Ramachandran plot, the determinant of the covariance matrix of the dihedral angles was calculated as:

$$\mathrm{detCMD} = \det\begin{pmatrix} \mathrm{Var}(\phi) & \mathrm{Cov}(\psi,\ \phi) \\ \mathrm{Cov}(\phi,\ \psi) & \mathrm{Var}(\psi) \end{pmatrix}$$

where $\mathrm{Var}(\phi)$ and $\mathrm{Var}(\psi)$ denote the variances of the $\phi$ and $\psi$ angles, respectively, and $\mathrm{Cov}(\phi,\ \psi)$ is covariance of these angles.

**MD data analysis: Ramachandran number**

The Ramachandran number $R$ for each residue was calculated using the formula (*11*):

$$R = \frac{\phi + \psi + 2\pi}{4\pi}$$

where $\phi$ and $\psi$ are expressed in radians. For each segment, $R$ values were calculated for all residues across 100 snapshots, resulting in 500 values per segment (5 residues × 100 snapshots). Residues with $R$ value between 0.475 and 0.575 were classified as extended structures, including β-strands, and those between 0.3 and 0.4 were classified as α-helix. Secondary structure proportions were determined from the distribution of $R$ values.

**MD data analysis: duration of extended structure and sidechain distance**

To assess the duration of extended structures, Ramachandran numbers were recorded for each residue at 1 ns interval, resulting in five time series (one per residue) per segment. For each residue, the longest continuous period during which the $R$ value remained between 0.475 and 0.575 was calculated. This process yielded five longest durations for extended structures per segment. The average of these five durations was used as the extended structure duration of the segment.

The sidechain distances were calculated as the distance between the centers of geometry of adjacent residue pairs (4 pairs in total: 1-2, 2-3, 3-4, 4-5) across 100 snapshots, yielding 400 distance values per segment (4 pairs × 100 snapshots). The center of geometry for each residue was calculated using MDAnalysis.

**3D-RISM calculation**

For the 3D-RISM calculations, 100 snapshots were taken from the MD trajectories every 1 ns and analyzed using RISMiCal program (*12*). All water molecules and ions were stripped from the snapshots and the 3D-RISM equation was solved by coupling with the Kovalenko-Hirata closure (*13*). TIP3P was used as water parameters because this type of 3 site models can be used in 3D-RISM calculations (*14*). The box size was $128^3$ Å$^3$ with 0.5 Å grid spacing.

**NMR sample preparation**

Uniformly [$^{13}$C, $^{15}$N]-labelled protein stocks were heated at 95 °C for 1 hour to ensure complete conversion to the monomeric state, then cooled to room temperature. The stocks were diluted to 500 µM in a buffer containing 12 % 1,6-HD, 50 mM MES, pH 6.5, 150 mM NaCl, 1 mM DTT, and 5 % $D_2O$. The samples were then heated at 95 °C for 10 min and transferred to 5 mm Shigemi tubes (SHIGEMI, Tokyo, Japan). Prior to NMR measurements, the samples were heated again in boiling water for 10 min and cooled to room temperature for 5 min. This procedure was performed to ensure sample homogeneity.

**NMR measurements and data analysis**

NMR experiments were conducted using an AVANCE III HD 600 MHz spectrometer equipped with a cryogenically cooled QCI-P probe (Bruker BioSpin GmbH, Billerica, MA, USA). Backbone resonance assignments were obtained using standard tripleresonance experiments, including hncacbgp3d, cbcaconhgp3d, hncogp3d, hncacogp3d, hncocagp3d and hncagp3d at 310 K. All spectra were acquired with Topspin 3.6.2 (Bruker) and processed using NMRPipe (*15*). Assignments were carried out using NMRFAM-Sparky and I-PINE web server (http://i-pine.nmrfam.wisc.edu/examples.html) (*16*, *17*) and were manually checked.

Relaxation measurements were performed using hsqct1etf3gpsi3d, hsqct2etf3gpsi3d and hsqcnoef3gpsi. For $T_1$ experiments, inversion recovery delays were set to 20, 100, 200, 400, 600, and 1,200 ms, with data points at 200 and 600 ms acquired in duplicate. For $T_2$ experiments, the relaxation delays were set to 16.96, 50.88, 84.8, 152.64, 203.52, 254.4, 407.04, 508.8 ms, with data points at 84.8 and 203.52 ms acquired in duplicate. $T_1$ and $T_2$ values were determined by fitting peak heights to a decaying exponential $h = A * \exp(-R * t)$ where $h$ is a peak height and $t$ is the delays for $T_1$ or $T_2$. Fitting was performed independently for two time series datasets including data points measured in duplicate, yielding two time constants $T$ (rate constant $R = 1/T$). The reported relaxation rates represent the averages of these two constants, with error bars indicating the standard deviation. Heteronuclear NOE experiments were performed when the saturation time was 3.0 s and the off-saturated time was 0 s.

The rotational correlation time was estimated from $T_1$ and $T_2$ as follows:

$$\tau_c \approx \frac{1}{4\pi\nu_N}\sqrt{6\frac{T_1}{T_2} - 7}$$

where $\nu_N$ is the $^{15}N$ resonance frequency (in Hz). In the above equation, $T_1$ and $T_2$ were substituted with their respective average values. This equation is reported at the NESG Wiki site (http://www.nmr2.buffalo.edu/nesg.wiki/NMR_determined_Rotational_correlation_time). The correlation time from the molecular weight was calculate by "Protein Correlation Time Calculators" (http://nickanthis.com/tools/tau.html), using a hydration layer of 3.2 Å. This estimation assumes a perfect spherical shape.

Secondary structure prediction was performed using TALOS-N dihedral angle calculation server (https://spin.niddk.nih.gov/bax/nmrserver/talosn/) (*18*). Backbone chemical shift data for HN, N, CA, CB, and CO were submitted to the server along with the amino acid sequence.

For the WT sample, the relaxation rates obtained in this study showed lower agreement with previously reported values (*2*). The prior measurement was performed using a Bruker 800 MHz spectrometer using a solution containing 10 % 1,6-HD, whereas the current study used a 600 MHz spectrometer with 12% 1,6-HD. To evaluate the impact of these differences, a WT sample containing 10% 1,6-HD was prepared and analyzed under the same conditions as described above using the 600 MHz spectrometer.

**CD measurement**

Samples were prepared at a protein concentration of 50 μM in PS buffer containing 12 % 1,6-HD, following the condensate sample preparation procedure. CD measurements were performed using a J-1500 CD Spectrometer (JASCO Corporation, Tokyo, Japan) and a quartz cell M20-B-3 with a path length of 0.005 cm (GL Sciences Inc., Tokyo, Japan).

Spectra were acquired at room temperature in the wavelength range of 190-250 nm. A solvent spectrum was measured as a baseline and subtracted from each sample spectrum. Spectra were smoothed using a fast Fourier transform in Spectra Manager software (JASCO).

The delta epsilon Δε (mdeg·$M^{-1}$·$cm^{-1}$) was calculated by the following formula:

$$[\Theta]_{MRE} = \frac{\Theta}{10 \cdot c \cdot N_r \cdot l}$$

$$\Delta\varepsilon = \frac{[\Theta]_{MRE}}{3298}$$

where Θ (mdeg) is the measured ellipticity, $c$ is the protein molar concentration (=0.00005 M), $N_r$ is the number of residues (=91), and $l$ is the path length (=0.005 cm).

Secondary structure estimation from the CD spectra was performed using the BeStSel web server (https://bestsel.elte.hu/) (*19*) with the following parameters: input unit, measured ellipticity; input range, 195–250 nm; protein concentration, 50 µM; number of residues, 91; path length, 0.005 cm.

**AF2 structure prediction**

To examine the relationship between the periodicity of F-r scores and β-sheet formation, we designed 30 new sequences for each lag using the same method described in the "Designing Lag-series IDP sequences" section. All generated sequences (180 in total) are listed in Dataset S1, sheet "newIDPsequence".

AlphaFold2 (AF2) was used to predict the tertiary structures of all lag-series IDPs, including newly generated sequences (*20*). A total of 186 sequences were input into AF2. Multiple sequence alignments (MSAs) were generated using the standard AF2 pipeline, and the predicted Lodal Distance Difference Test (pLDDT) score was used to rank model confidence. Five models were generated for each sequence, and Amber relaxation was performed on all models.

Secondary structure assignments were performed using the DSSP (Define Secondary Structure of Proteins) program (*21*). To calculate secondary structure (SS) probability for each residue, the number of models predicting β-sheet or α-helix conformation at that residue was counted and divided by 5, yielding SS probability by the following formula:

$$\text{SS probability}(i) = N_i(\beta\text{-sheet or }\alpha\text{-helix})/5$$

where $i$ represents a residue number, and $N_i$ represents the number of models predicting β-sheet or α-helix conformation at residue number $i$.

To quantify secondary structures content for each sequence, the sum of SS probabilities, $E(\beta\text{-sheet or }\alpha\text{-helix})$, was calculated as the expected number of residues forming β-sheet or α-helix conformations per sequence by the following formula:

$$E(\beta\text{-sheet or }\alpha\text{-helix}) = \sum_{i=1}^{N} \text{SS probability}(i)$$

where $i$ represents a residue number, and $N$ represents the total number of residues in lag-series IDPs, and SS $\mathrm{probability}(i)$ represents the probability of forming β-sheet or α-helix conformations at residue number $i$. We refer to this value as the predicted β-sheet content.

Correlation between β-sheet probability and F-r scores was then analyzed. Initially, we attempted to calculate correlation coefficients for each individual sequence. However, this approach did not work appropriately, as some sequences lacked predicted β-sheet regions, making correlation calculations infeasible. This limitation introduced inconsistencies in the number of analyzable sequences per group, potentially biasing the results. To address this issue, we developed an alternative approach. Rather than analyzing individual sequences, we combined the β-sheet probabilities across all sequences within each group. The combined β-sheet probability was calculated using the following formula:

$$\mathrm{SSprob\ comb} = \left(\mathrm{SSprob}_1(i), \mathrm{SSprob}_2(i), \cdots, \mathrm{SSprob}_\mathrm{N}(i)\right)$$

where $\mathrm{SSprob}_\mathrm{k}(i)$ denotes the β-sheet probability at residue number $i$ for the k-th sequence in the group, and N is the number of sequences within the group. For each sequence, the per-residue probability $\mathrm{SSprob}_\mathrm{k}(i)$ was defined as:

$$\mathrm{SSprob}_\mathrm{k}(i) = \left(\mathrm{SSprob}_\mathrm{k}(1), \mathrm{SSprob}_\mathrm{k}(2), \cdots, \mathrm{SSprob}_\mathrm{k}(n)\right)$$

where $n$ represents the total number of residues.

Similarly, we combined the F-r scores of all sequences within each group. The correlation coefficients were then calculated between the combined β-sheet probabilities and the combined F-r scores across 50 different AA-index values. This revised approach enabled the inclusion of all sequences while minimizing group-to-group bias, providing a more robust analysis of the relationship between β-sheet propensity and F-r scores across different groups.

**Statistical analysis**

All *p*-values were two sided and the significance level was set to be $\alpha = 0.05$ beforehand with the analyses. All statistical analyses were performed with EZR (Saitama Medical Center, Jichi Medical University, Saitama, Japan; Kanda, 2013) (*22*), which is a graphical user interface for R (The R Foundation for Statistical Computing, Vienna, Austria). More precisely, it is a modified version of R commander designed to add statistical functions frequently used in biostatistics.

### Supplementary Figures

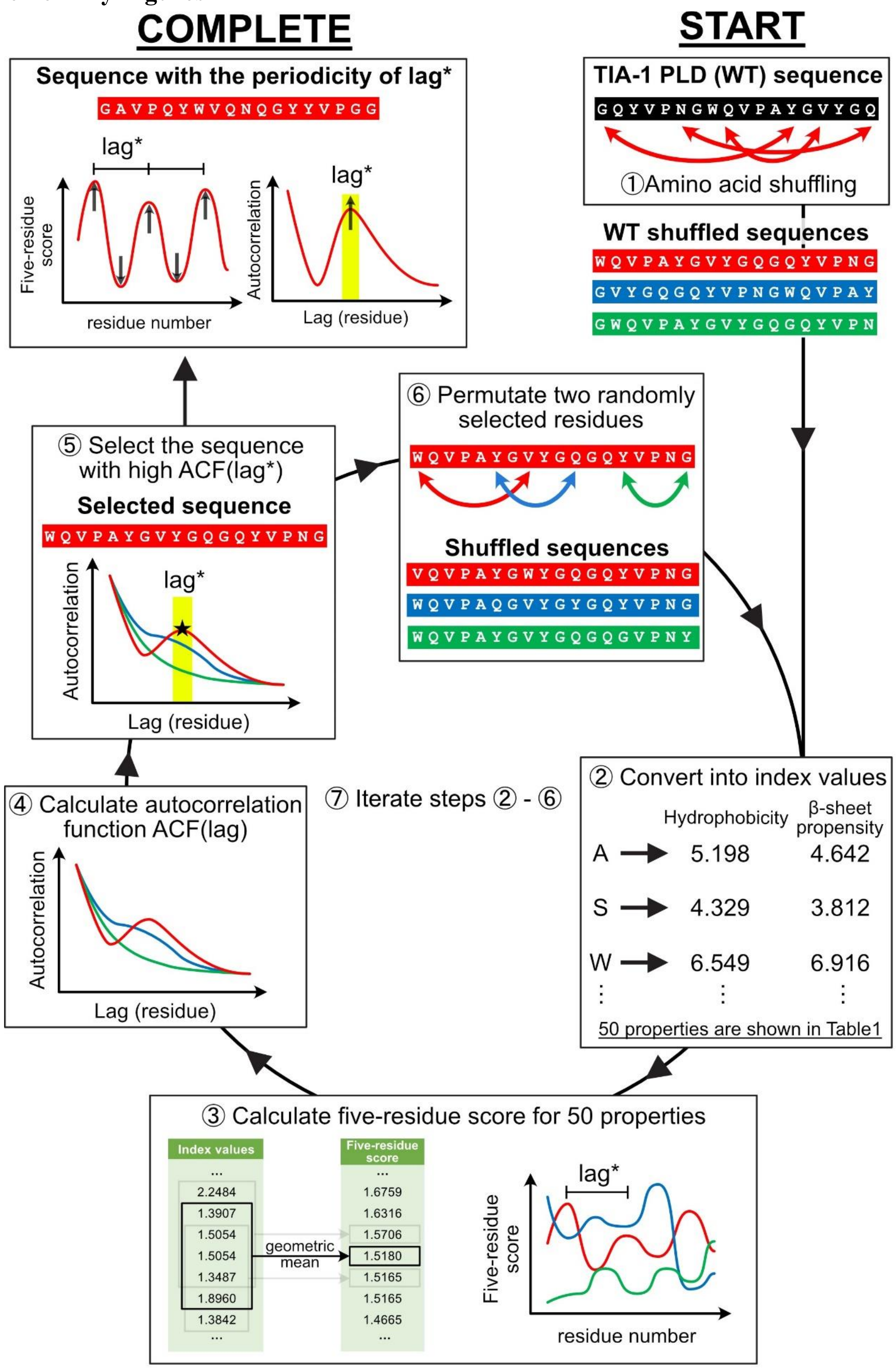

**Fig. S1. Schematic procedure for designing lag-series IDPs.**

The aim of this procedure is to obtain IDP sequences with a periodicity of F-r scores. Iterative cycles shown in the centre progressively select IDP sequences with stronger periodicity of the F-r scores at a lag*. In the final step, the sequence exhibiting the highest periodicity is selected. The following is an explanation of each step.

1. The wild-type (WT) TIA-1 PLD sequence was randomly shuffled to generate a pool of sequences (WT shuffled sequences).
2. Each sequence was converted into 50 physicochemical indices, such as hydrophobicity or β-sheet propensity (Dataset S1, sheet ”indexvalues”).
3. Five-residue scores (F-r scores) were calculated as the geometric mean of the indices for every five residues. We hypothesize that F-r scores reflect conformational biases: residues with high F-r scores possess strong bias and slow dynamics, while residues with low F-r scores possess weak bias and fast dynamics.
4. Autocorrelation functions, ACF(lag), of the F-r scores were calculated to quantify periodicity.
5. For a target periodicity lag* (10, 20, 30, 40, and 50 residues), the sequence with the highest ACF(lag*) value was selected from the WT shuffled sequences (Selected sequence).
6. For the selected sequences, two residues were randomly swapped to generate a new pool of shuffled sequences (Shuffled sequences).
7. Steps 2-6 were iterated until ACF(lag*) converged.
8. To obtain a sequence with low periodicity, referred to the non-periodic sequence (NP), the sequence with minimal summed ACF(lag) and subjected to the same iterative shuffling.

Using this method, we generated artificial sequences named L10, L20, L30, L40, and L50, each corresponding to its respective lag* value, as well as NP, which lacked periodicity.

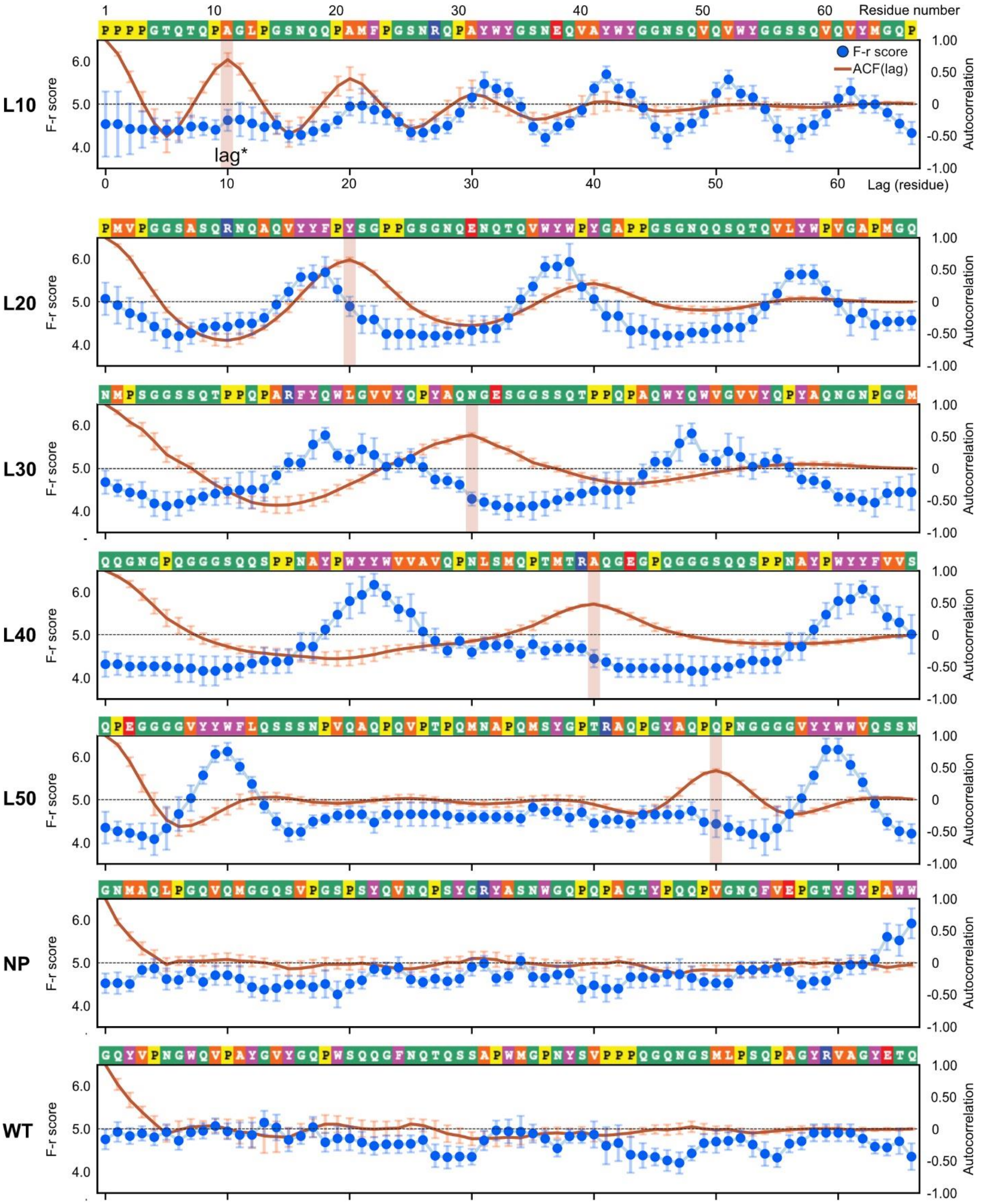


**Fig. S2. F-r scores and autocorrelation functions ACF(lag) of lag-series IDPs.**
Each panel shows the F-r scores and autocorrelation functions ACF(lag). Blue data points represent the mean value of the F-r scores, with error bars indicating the standard deviation of the F-r scores. Red solid lines represent the mean value of ACF(lag) calculated from 50 types of

the F-r scores, with error bars indicating the standard deviation of the ACF(lag) values. Amino acid sequences are shown above the graphs, and background colors for each residue indicate the amino acid types: green for polar (G, T, Q, S, N), orange for hydrophobic (A, L, M, V), magenta for aromatic (F, W, Y), yellow for proline (P), red for acidic (E), and blue for basic (R) residues.

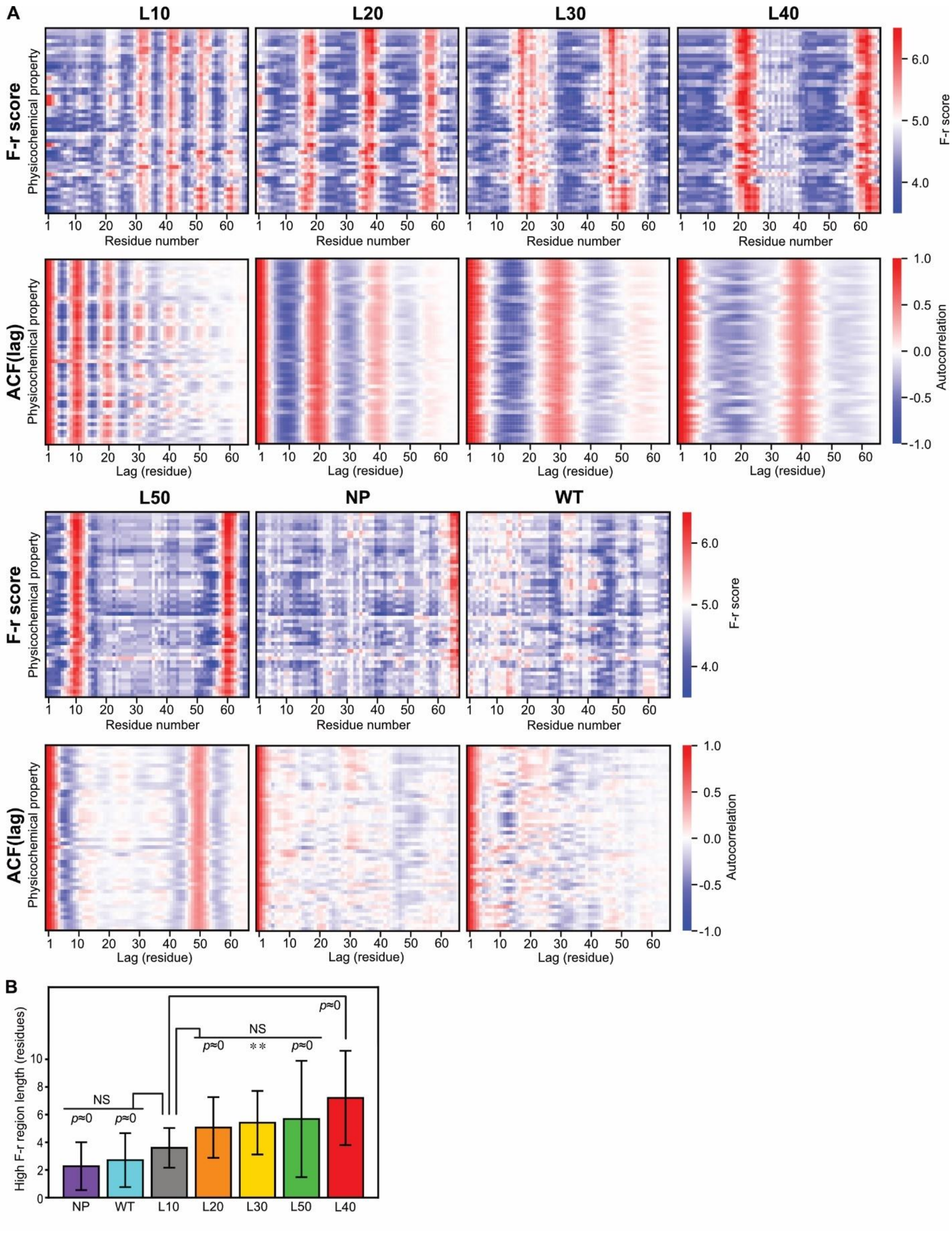


**Fig. S3. F-r scores and ACF(lag) for lag-series IDPs.**
(**A**) All F-r scores and ACF(lag) values of lag-series IDPs are shown as heatmaps. Heatmaps display the F-r scores or ACF(lag) values calculated from 50 types of index values representing

physicochemical properties for each amino acid. Along the vertical axis, each row represents the F-r scores or ACF(lag) values corresponding to each physicochemical property. The vertical axis represents the residue number in the F-r scores, and the lag value in ACF(lag), respectively. The color bar indicates the magnitude of the F-r scores or ACF(lag) values, respectively. F-r score values are shown in the Dataset S1.
(**B**) The lengths of high F-r score regions in lag-series IDPs. Contiguous regions with the F-r scores higher than 5.0 were counted across all 50 types of the F-r scores. The threshold of 5.0 was used to distinguish between high and low F-r scores, as the F-r scores were calculated from index values normalized to an average. Bar plots represent mean ± SD. Steel Dwass' method was used for multiple comparisons. Asterisks represents adjusted *p*-value vs. L10, **: $p < 5.0 \times 10^{-3}$ and NS: no significant difference. $p \approx 0$ represents $p < 1.0 \times 10^{-7}$.

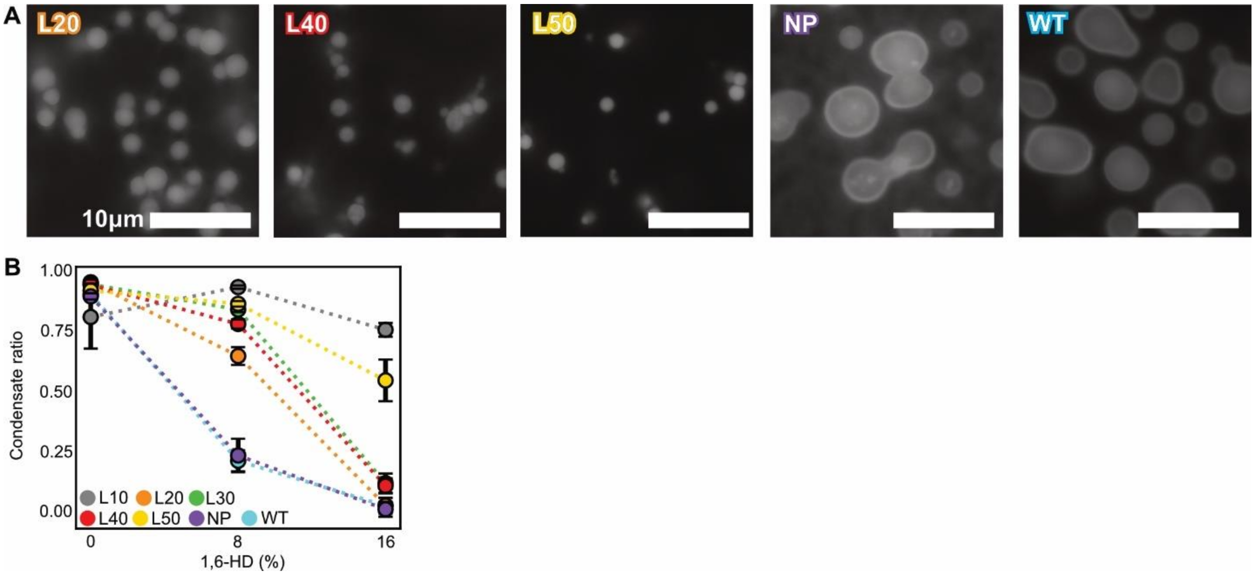


**Fig. S4. Fluorescence images and condensate ratio of lag-series IDPs.**
(**A**) Representative fluorescence microscopy images of condensates formed by lag-series IDPs. For accurate measurements of condensate sizes, thioflavin-T (ThT) was added to visualize the condensates in fluorescence microscopy. ThT fluorescence reflects the viscosity of the condensates, not amyloid fibrils, as images were acquired within 20 minutes of mixing, before fibril growth occurs. Scale bar, 10 μm.
(**B**) Condensate ratios of lag-series IDPs. Samples were prepared in the same buffer as that in A with 0, 8, or 16 % 1,6-HD. Data points represent mean ± SD (N = 3). One-way ANOVA showed a significant difference at 8 % 1,6-HD ($p = 8.32\times10^{-13}$).

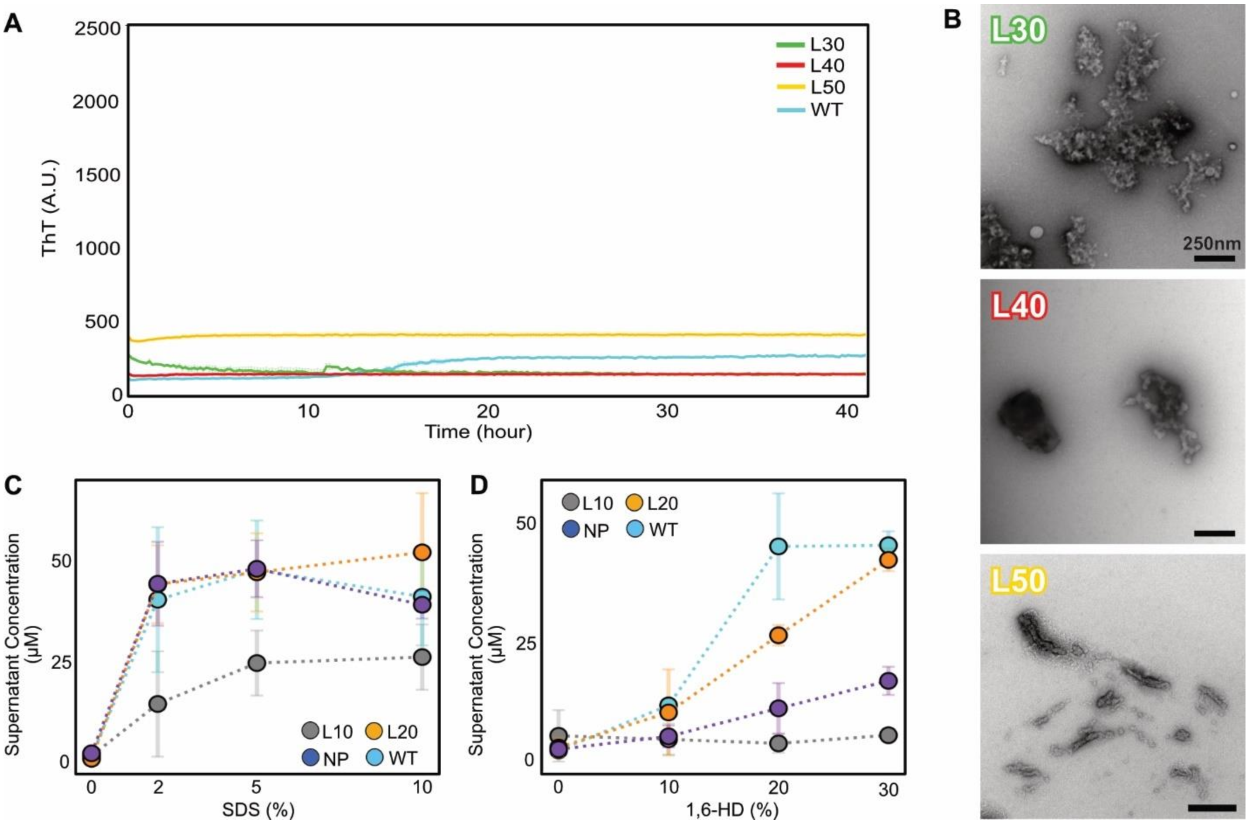


**Fig. S5. Amyloid fibril formation and condensate stability of lag-series IDPs including WT.**
(**A**) ThT assay of L30, L40, L50, and WT. Excitation 445 nm, emission 485 nm. Curves show mean ± SD (N = 3).
(**B**) TEM images of L30, L40, L50, and WT. Samples were prepared by incubating the same solution as that in A at room temperature for 72 hours. Scale bar, 250 nm.
(**C**) SDS resistance of L10, L20, NP, and WT condensates. Condensate samples of L10, L20, and NP were prepared using the same method as the negative staining TEM. Subsequently, SDS was added to the samples at concentrations corresponding to those shown in the graph, and the samples were incubated for 24 hours at room temperature. The solutions were centrifuged and the protein concentration in the supernatants was measured. Data points represent mean ± SD (N = 3). One-way ANOVA showed a significant difference at 5 % SDS ($p$ = 0.0409).
(**D**) 1,6-HD resistance of L10, L20, NP, and WT condensates. The same method used for the SDS resistance assay was employed, with 1,6-HD instead of using SDS. Data points represent mean ± SD (N = 3). One-way showed a significant difference between constructs at 20 % 1,6-HD ($p = 2.03 \times 10^{-4}$) and 30 % 1,6-HD ($p = 8.14 \times 10^{-8}$).

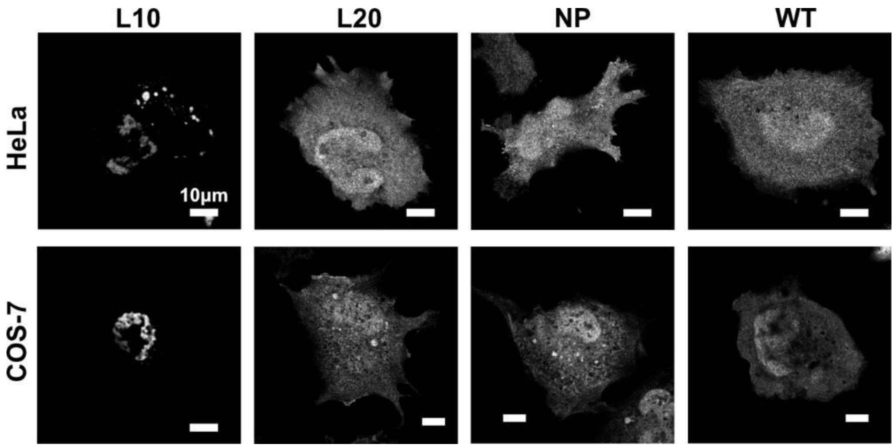


**Fig. S6. Intracellular condensate of lag-series IDPs.**
Super resolution fluorescence microscopy images of HeLa cells and COS-7 cells expressing L10, L20, NP and WT fused with EYFP. Scale bar, 10 μm.

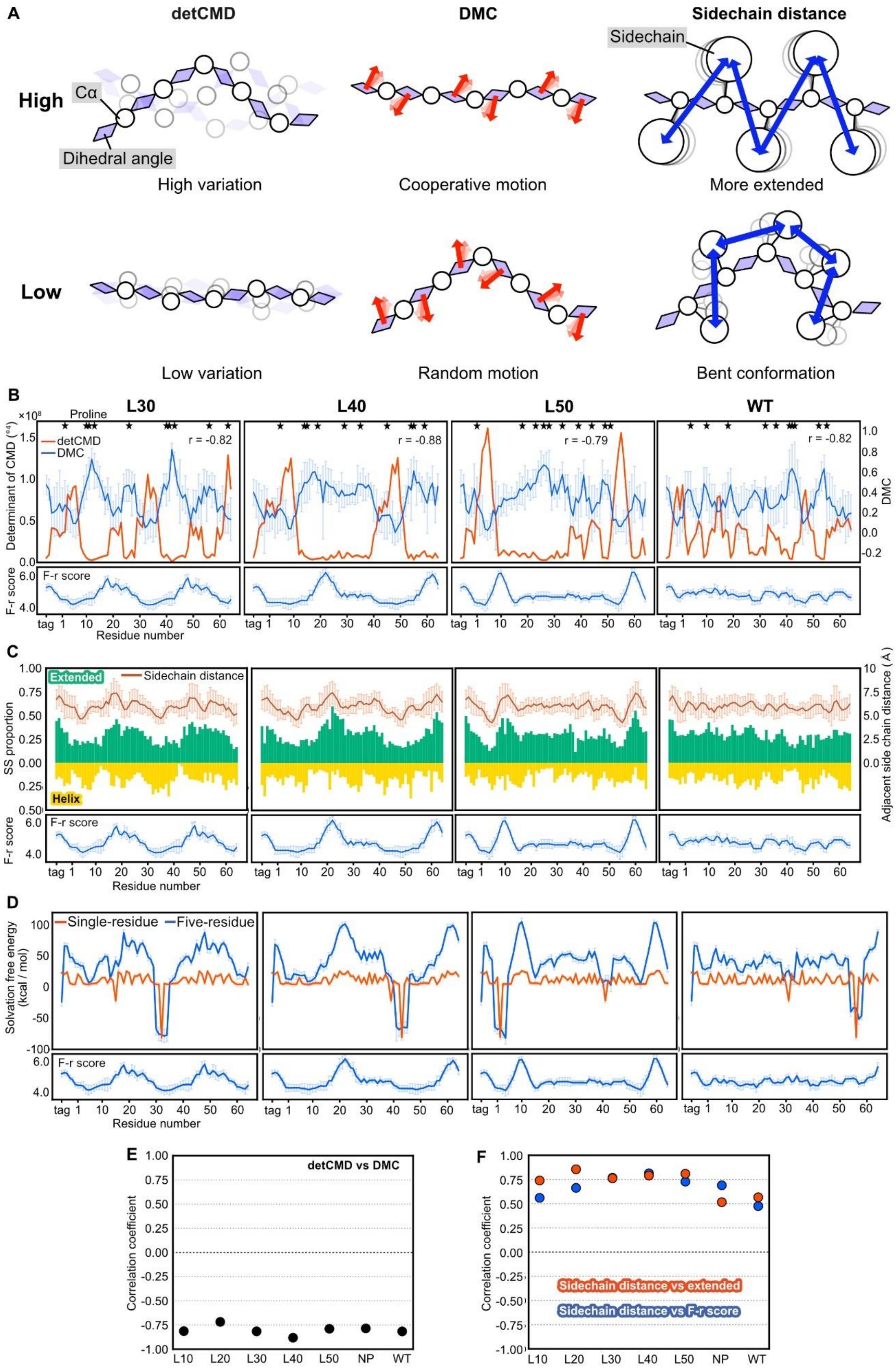

A
detCMD
DMC
Sidechain distance
High
Cα
Dihedral angle
Sidechain
High variation
Cooperative motion
More extended
Low
Low variation
Random motion
Bent conformation
B
L30
L40
L50
WT
Proline
detCMD
DMC
r = -0.82
r = -0.88
r = -0.79
r = -0.82
Determinant of CMD
F-r score
Residue number
C
Extended
Helix
Sidechain distance
SS proportion
Adjacent side chain distance (Å)
F-r score
Residue number
D
Single-residue
Five-residue
Solvation free energy (kcal / mol)
F-r score
Residue number
E
detCMD vs DMC
Correlation coefficient
L10
L20
L30
L40
L50
NP
WT
F
Sidechain distance vs extended
Sidechain distance vs F-r score
Correlation coefficient

**Fig. S7. MD simulations for five-residue segments of L30, L40, L50, and WT.**
(**A**) Schematic diagram of determinant of the covariance matrix of the backbone dihedral angles (detCMD), dihedral angle motion correlation (DMC), and sidechain distance. Each parameter was calculated for five-residue segments. Multiple overlapping structures reflect the structural ensemble. High detCMD indicates greater structural variation, while low detCMD indicates restricted conformations. For DMC, red arrows represent dihedral motions, where high DMC reflects cooperative motion and low DMC reflects random motion. For sidechain distance, longer distances between sidechains lead to extended backbones, while shorter distances favor bent conformations.
(**B**) DMC and detCMD. The lower panel shows the corresponding F-r scores. Proline positions are marked by stars. The correlation coefficients between DMC and the detCMD are shown as *r*.
(**C**) Secondary structure (SS) proportions and sidechain distances of five-residue segments. The lower panel shows the corresponding F-r scores. Green and yellow bars represent extended and α-helical structures, respectively. SS proportions were calculated from the distributions of Ramachandran numbers (Fig. S9). Lines represent the mean values of all sidechain distances of four adjacent residue pairs, with error bars indicating SD.
(**D**) Solvation free energy (SFE) is shown in the upper panel with F-r scores shown in the lower. Blue lines represent the mean SFE values calculated for five-residue segments, with error bars indicating SD. Red lines represent the mean SFE values calculated for single residues, with error bars indicating SD.
(**E**) Correlation coefficients between detCMD and DMC. Data points represent correlations between DMC detCMD.
(**F**) Correlation coefficients between F-r scores or extended structure proportions and sidechain distances. Blue data points represent those between sidechain distances and F-r scores; red data points represent correlations between sidechain distances and extended structure proportions.

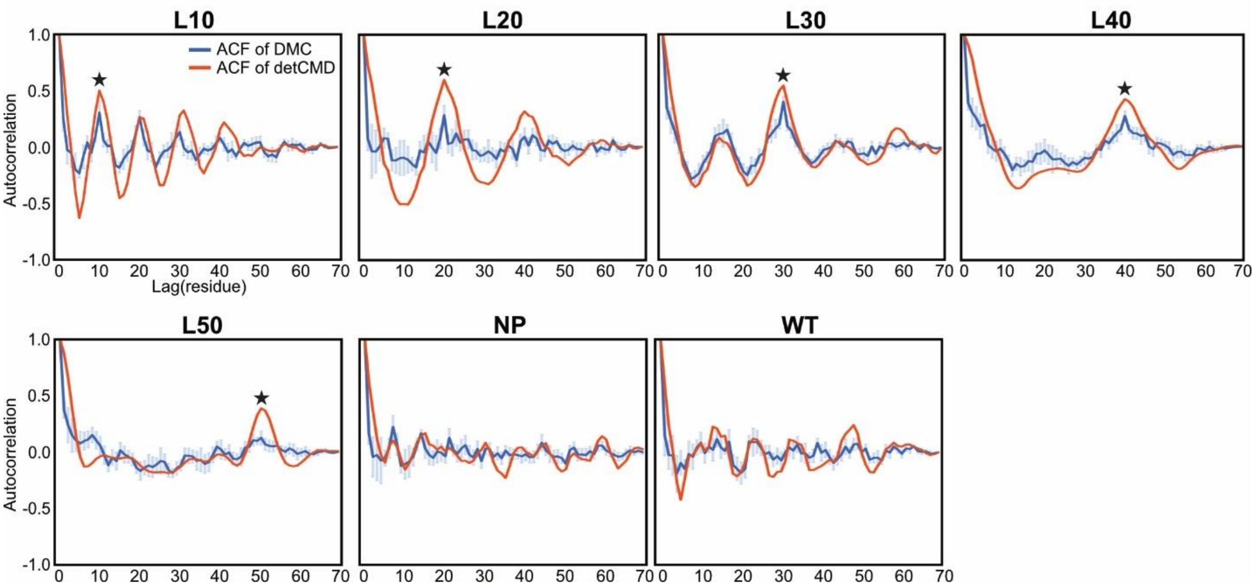


**Fig. S8. Autocorrelation functions (ACF) of DMC and detCMD of all lag-series IDPs.** Blue lines represent the mean value of ACF calculated from 10 values of DMC, with error bars indicating the standard deviation of the ACF values. Red lines represent the ACF calculated from detCMD. Data used for ACF calculations are shown in Fig. 4A and Fig. S7B. Stars in each panel except for NP and WT indicate the position of lag* for each lag-series IDP.

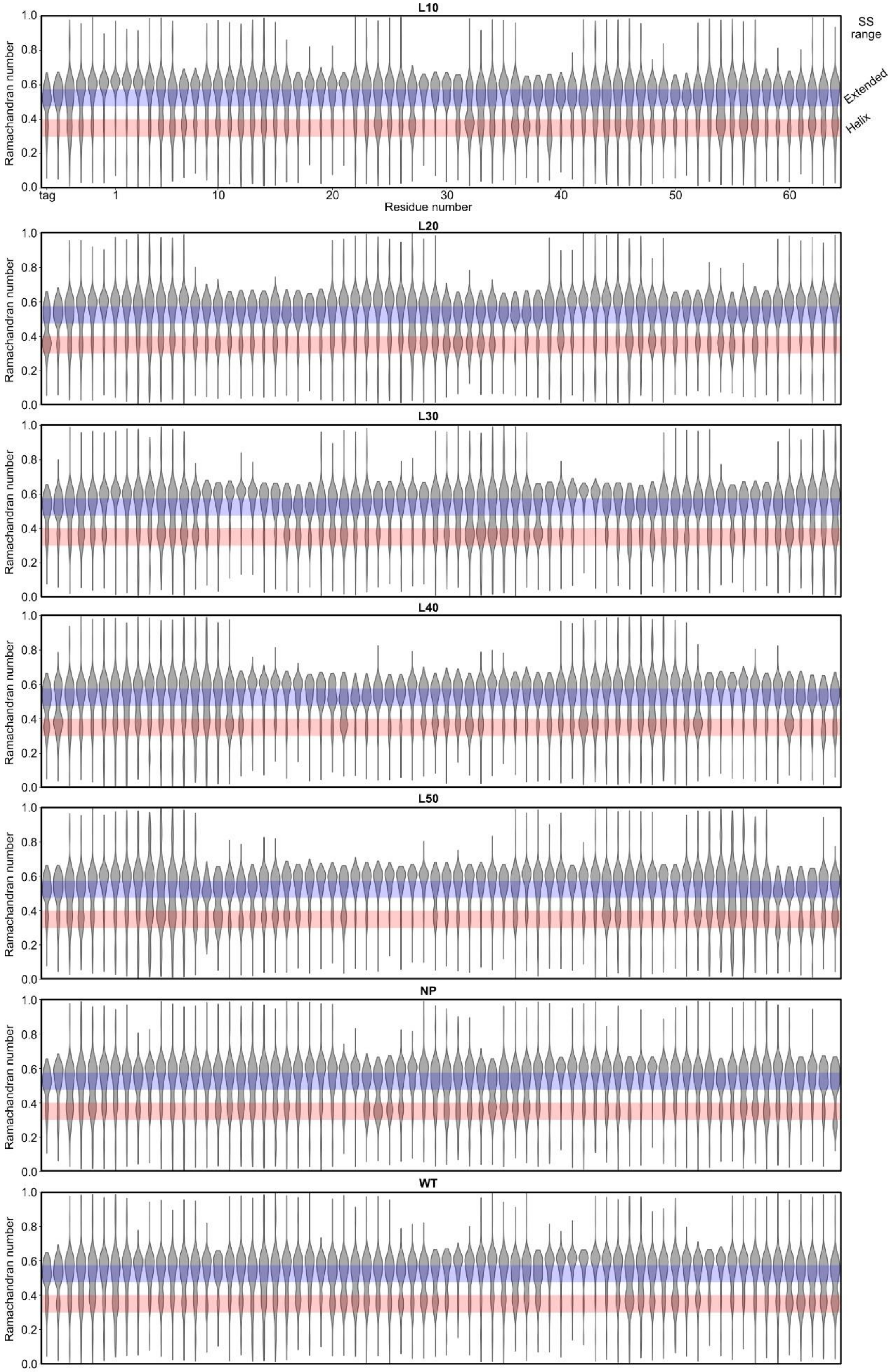

L10
L20
L30
L40
L50
NP
WT
Ramachandran number
Residue number
tag
SS range
Extended
Helix

**Fig. S9. The Ramachandran number distributions of lag-series IDPs.**
Violin plots of the Ramachandran numbers calculated from the MD trajectories of lag-series IDPs. The edges of each violin plot are the maximum and minimum values. Blue shaded area represents the range of Ramachandran numbers indicating extended structures, and red represents the α-helix. These areas are shown as SS range on the right. The populations included in each area are shown in Fig. 4B and S7C.

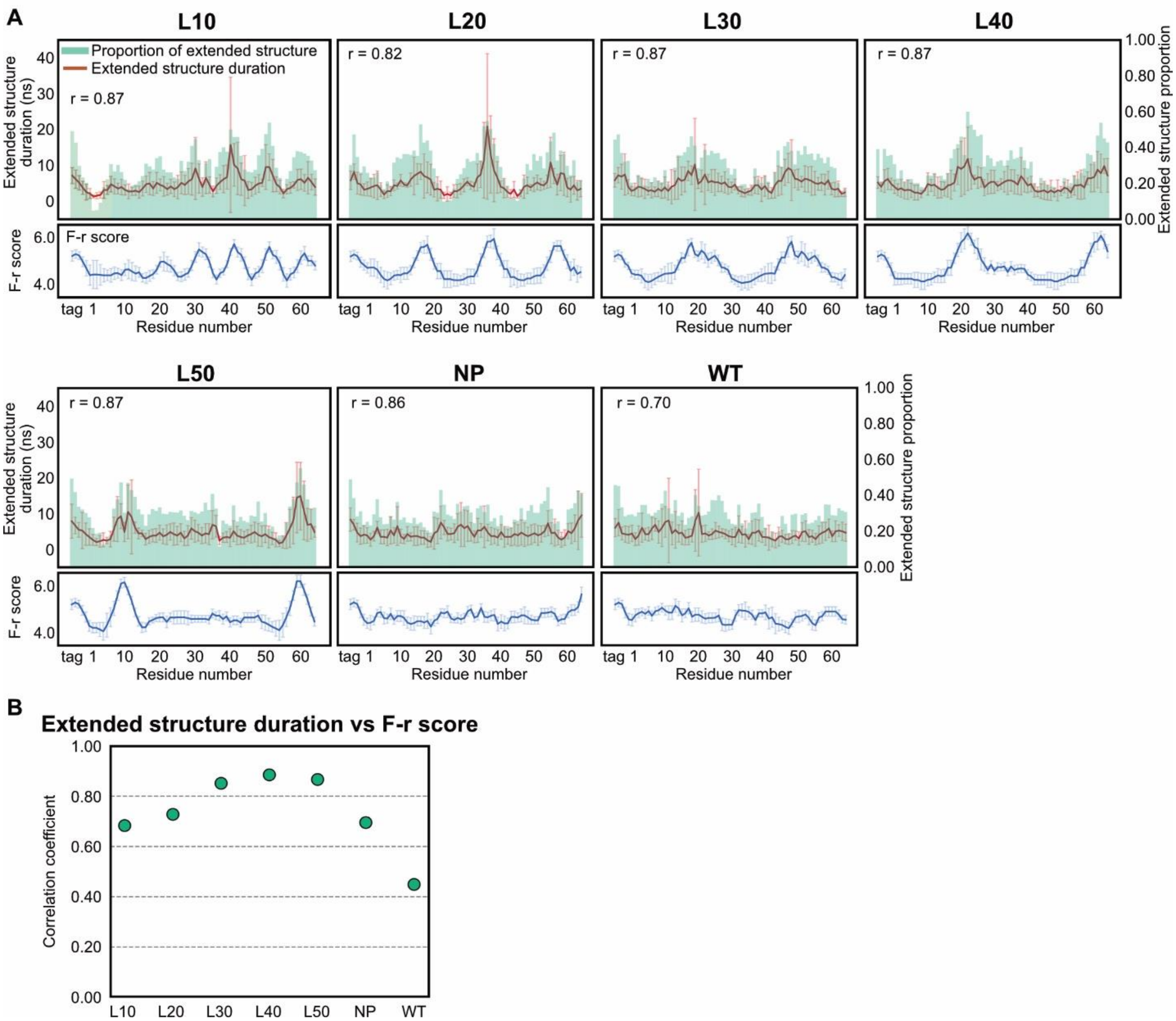


**Fig. S10. The extended structure durations of lag-series IDPs.**

(**A**) Proportions and durations of extended structures. The proportions and durations of extended structures are shown in upper panels, and the F-r scores are shown in lower panels. Extended structures correspond to the structures with the Ramachandran number, $R$, between 0.475 and 0.575. Green bars represent the proportions of extended structures. The durations of extended structures (extended structure duration) were calculated as the longest consecutive time to maintain the extended structures from the $R$ values for each residue per segment. Red lines represent the mean of five values of extended structure durations per segment, with error bars indicating the standard deviations. The correlation coefficients between the mean values of extended structure durations and the proportions of extended structures are shown as $r$ in each panel. The F-r scores are shown in the same way as in Fig. 1B.

(**B**) Correlation coefficients between extended structure duration and F-r score. Data points represent the correlation coefficients between the mean values of extended structure durations and F-r scores

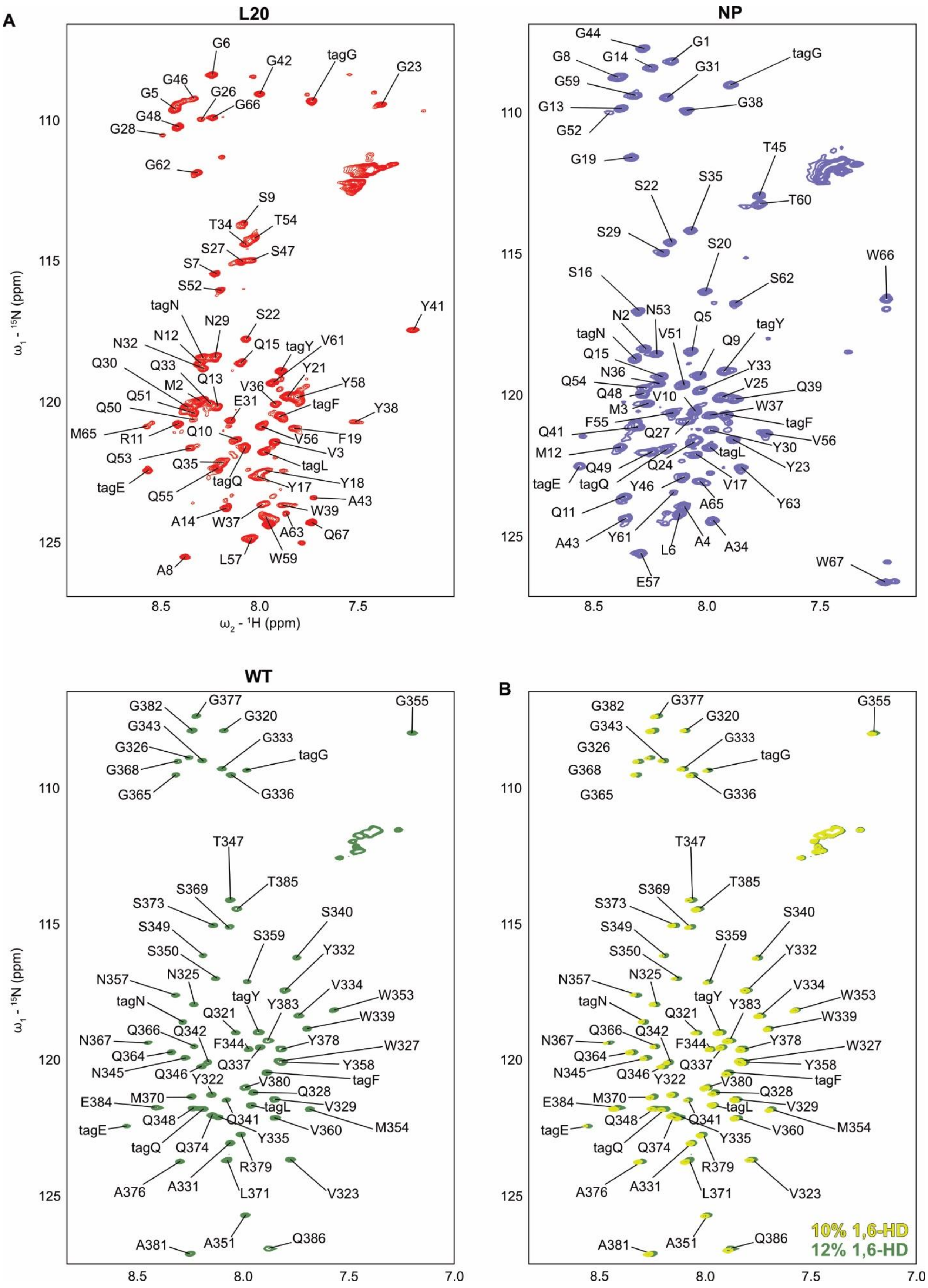

A
L20
NP
WT
B
G6
G42
tagG
G23
G46
G5
G26
G66
G48
G28
G62
S9
T34
T54
S27
S47
S7
S52
tagN
N29
S22
Y41
N32
N12
Q15
tagY
V61
Q30
Q33
Q13
Y21
M2
V36
Y58
Q51
E31
tagF
Q50
Y38
M65
R11
Q10
V56
F19
V3
Q53
Q35
tagL
tagE
tagQ
Y17
Y18
Q55
A43
A14
W37
W39
A63
Q67
L57
W59
A8
G44
G1
G8
G14
G31
G59
G13
G38
G52
G19
T45
S35
T60
S22
S29
S20
W66
S16
S62
N2
N53
Q5
tagN
V51
Q9
Q15
N36
Y33
Q54
Q48
V25
Q39
V10
M3
W37
F55
tagF
Q41
Q27
V56
Y30
M12
Q24
tagL
Q49
Y23
tagE
tagQ
Y46
V17
Y63
A65
Q11
A43
Y61
A4
A34
L6
W67
E57
G382
G377
G355
G320
G343
G333
G326
tagG
G368
G365
G336
T347
T385
S369
S373
S340
S349
S359
Y332
S350
N357
N325
V334
W353
tagN
tagY
Q321
Y383
W339
Q366
Q342
N367
F344
Y378
W327
Q364
Q337
Y358
N345
Q346
Y322
tagF
V380
Q328
E384
M370
V329
tagL
Q348
Q341
M354
tagE
V360
Y335
tagQ
Q374
R379
A376
A331
L371
V323
A351
Q386
A381
10% 1,6-HD
12% 1,6-HD
$\omega_1$ - $^{15}$N (ppm)
$\omega_2$ - $^{1}$H (ppm)
110
115
120
125
8.5
8.0
7.5
7.0

**Fig. S11. Backbone assignments of L20, NP, and WT in $^{1}H$-$^{15}N$ HSQC spectra.**
(**A**) The $^{1}H$-$^{15}N$ HSQC spectra of L20, NP and WT with their assignments. Residue numbers of WT are the same as our previous measurement (*2*).
(**B**) The $^{1}H$-$^{15}N$ HSQC spectra of WT in a buffer containing 10 % 1,6-HD and 12 % 1,6-HD, respectively. The $^{1}H$-$^{15}N$ HSQC spectrum of WT in a buffer containing 10 % 1,6-HD is shown in yellow, while that with 12% 1,6-HD is shown in green.

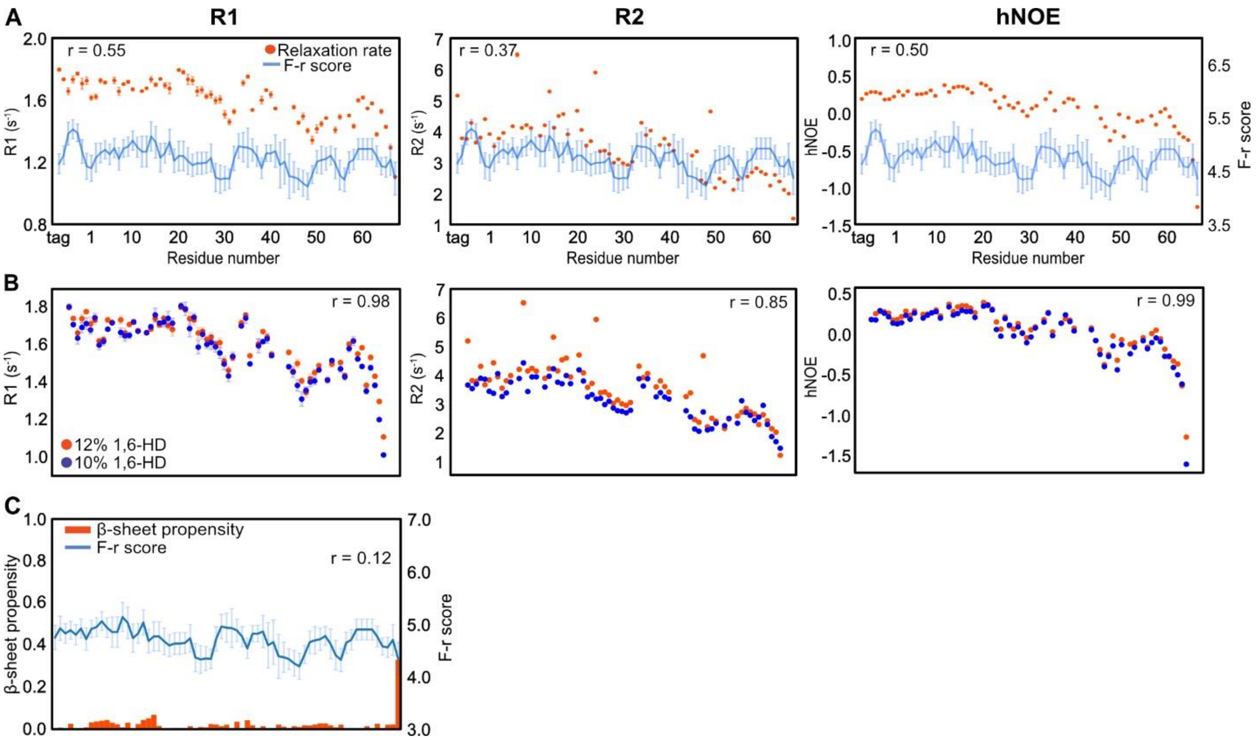


**Fig. S12. Relaxation rates and β-sheet propensity of WT.**

(**A**) $R_1$, $R_2$ and hNOE of WT. Relaxation rates were estimated from two time series datasets, and data points represent mean ± SD. The F-r scores are shown in the same way as in Fig. 1B. The correlation coefficients between the relaxation rates and the mean values of the F-r scores are shown as *r* in each panel.

**(B)** Comparison of the relaxation rates of WT in a buffer containing 10 % and 12 % 1,6-HD. Relaxation rates of $R_1$, $R_2$, and hNOE for WT samples in a buffer containing 10 % (blue) and 12 % 1,6-HD (red). Relaxation rates were estimated from two time series datasets, and data points represent mean ± SD. The F-r scores are shown in the same way as in Fig. 1B. The correlation coefficients between the relaxation rates of the 10 % and 12 % 1,6-HD samples are shown as *r* in the upper panels, respectively.

(**C**) β-sheet propensity of WT. Bar plots represent β-sheet propensity estimated from backbone chemical shifts, and the F-r scores are shown in the same way as in Fig. 1B. The correlation coefficients between the β-sheet propensity and the mean values of the F-r scores are shown as *r* in each panel. For α-helix propensities, see Fig. S13.

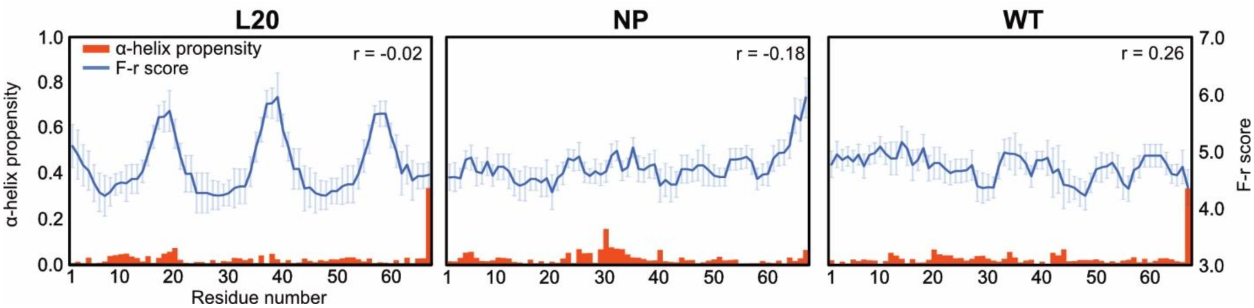


**Fig. S13. α-helix propensities of L20, NP, and WT.**

α-helix propensity of L20, NP, and WT. Bar plots represent α-helix propensity estimated from backbone chemical shifts, and the F-r scores are shown in the same way as in Fig. 1B. The correlation coefficients between the α-helix propensity and the mean values of the F-r scores are shown as *r* in each panel.

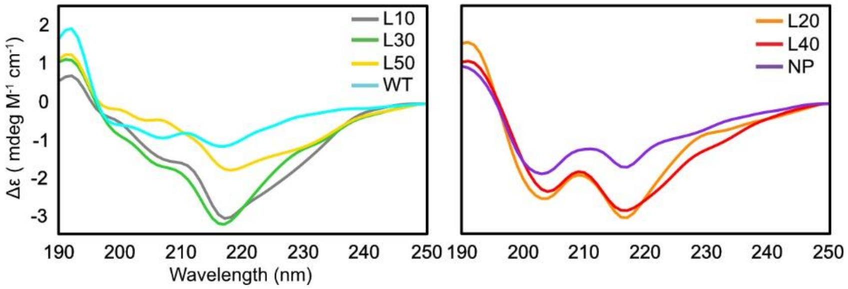


**Fig. S14. Circular dichroism (CD) spectra of lag-series IDPs.**
The delta epsilon (Δε) calculated from the measured ellipticities are plotted for each sample. For clarity, the spectra are divided into two panels: L10, L30, L50, and WT (left), and L20, L40, and NP (right), to accommodate differences in spectral features and improve visual separation of overlapping traces.

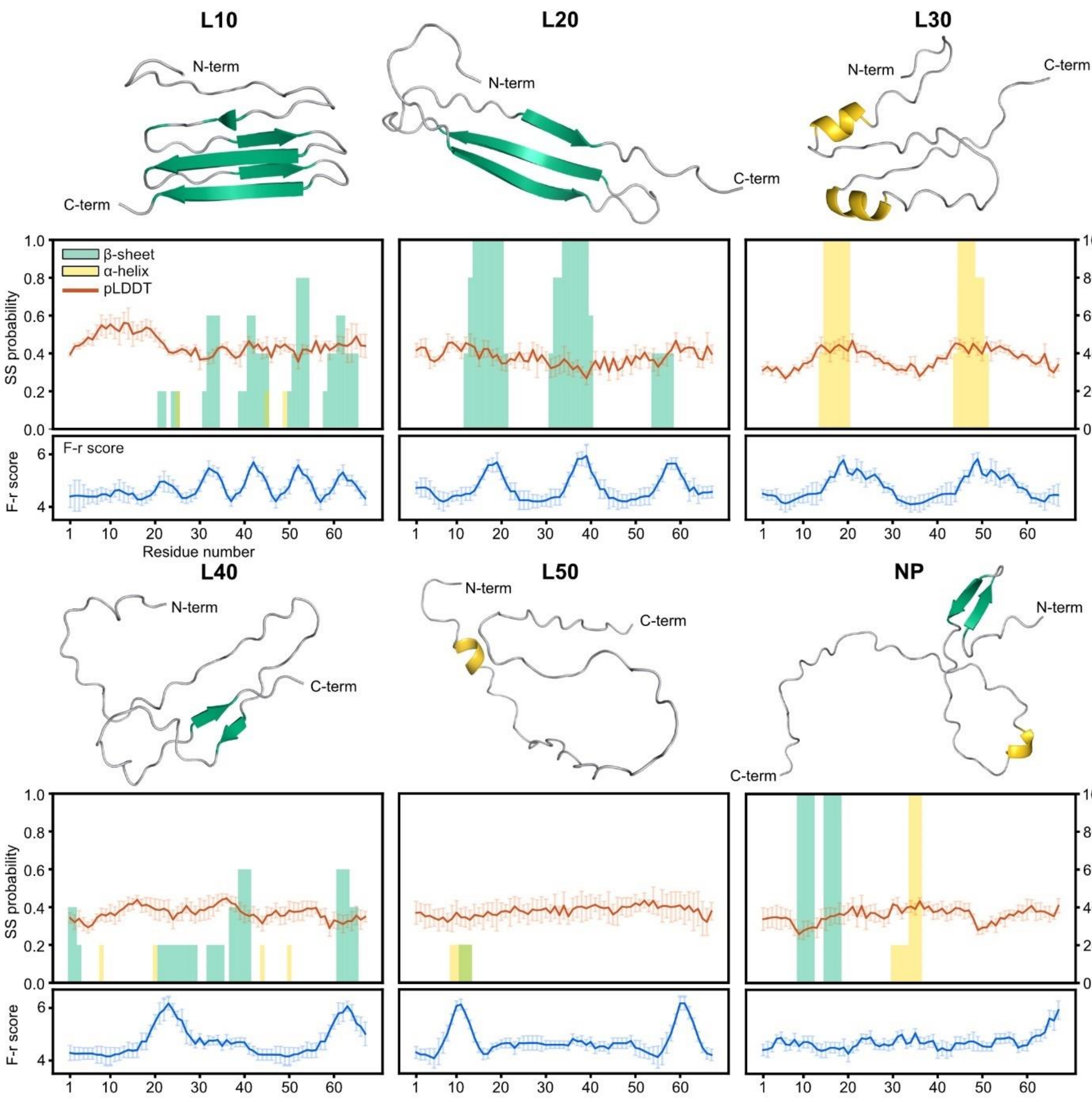


**Fig. S15. AlphaFold2 (AF2) predictions for lag-series IDPs.**

For each IDP, a representative structure from the top five ranked models is shown at the top. In these structures, β-sheets and α-helices are colored green and yellow, respectively. Secondary structure (SS) probabilities and pLDDT scores are shown in the panels below each structure. SS probabilities shown in bar plots were calculated for each residue as the proportion of models exhibiting β-sheet or α-helix conformations among the top five ranked models at each residue. The pLDDT plots represent the mean values across all five models, with error bars indicating standard deviations. The F-r scores are shown in the same way as in Fig. 1B.

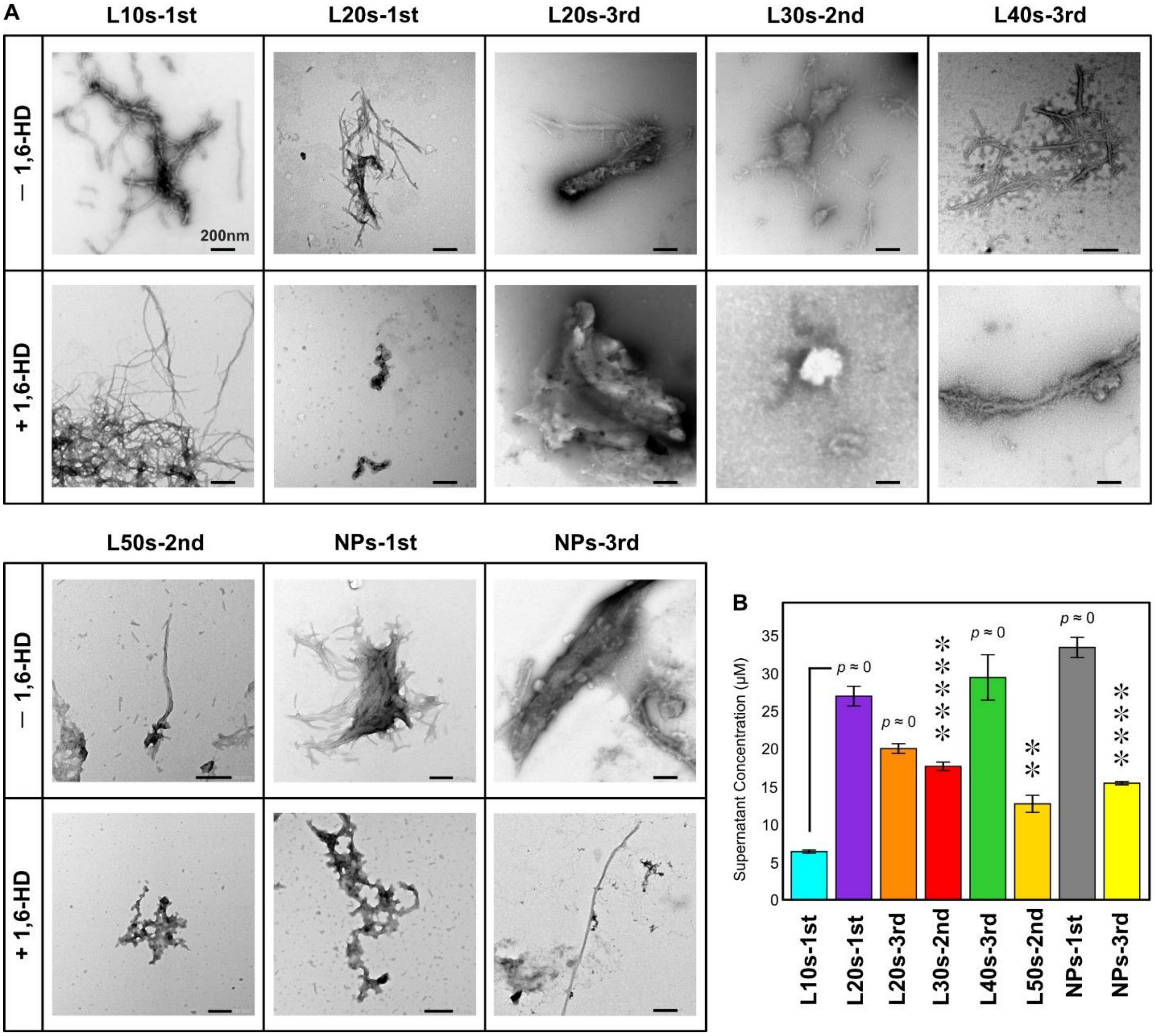


**Fig. S16. Amyloid fibrils and condensate stability of newly designed lag-series IDPs.**
(**A**) Negative-staining TEM images of condensate samples before and after 1,6-HD treatment. Condensate samples were prepared by incubating 25 μM IDPs at room temperature for 72 hours. Images labeled as –1,6-HD were taken prior to treatment. For +1,6-HD, 1,6-HD was added to a final concentration of 20%, followed by incubation for 24 hours at room temperature. Scale bar, 200 nm.
**(B)** 1,6-HD resistance of newly designed lag-series IDP condensates. Condensate samples were prepared under the same conditions as for TEM, using 50 μM IDPs. After 72 hours of incubation, 1,6-HD was added to a final concentration of 30%, and samples were incubated for 24 hours at room temperature. The solutions were centrifuged and the protein concentration in the supernatants was measured. Data points represent mean ± SD (N = 3). Tukey's method was used for multiple comparisons. Asterisks represents adjusted *p*-value vs. L10s-1st, **: $p < 5.0 \times 10^{-3}$ , ****: $p < 5.0 \times 10^{-5}$, *****: $p < 5.0 \times 10^{-6}$, $p \approx 0$ represents $p < 1.0 \times 10^{-7}$, and NS: no significant difference.

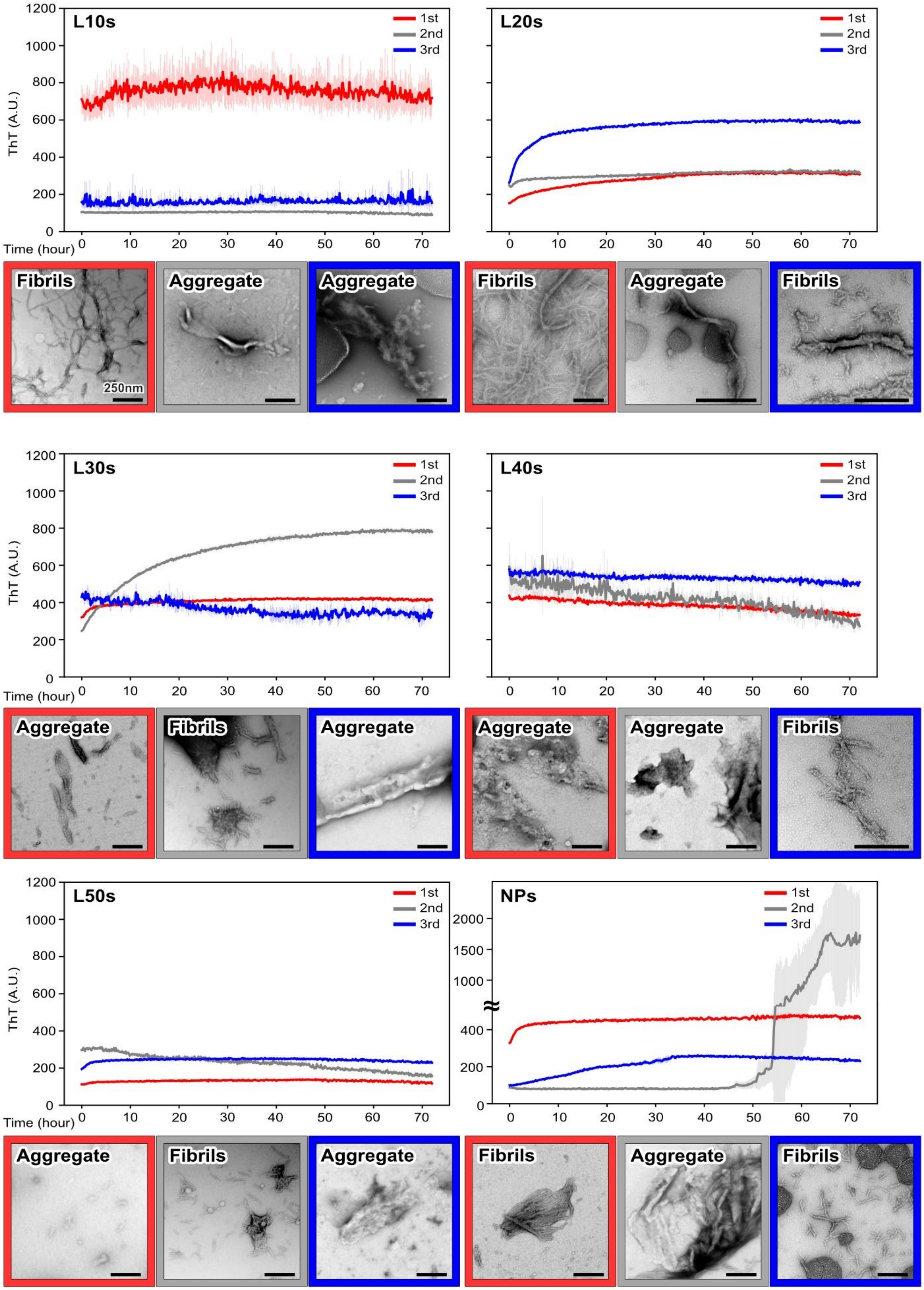

L10s
L20s
L30s
L40s
L50s
NPs
1st
2nd
3rd
ThT (A.U.)
Time (hour)
Fibrils
Aggregate
250nm

**Fig. S17. Amyloid fibril formation of newly designed lag-series IDPs.**

ThT assay and negative staining TEM images are shown in each panel for three representative sequences with highest periodicity score. For ThT assay, excitation 445 nm, emission 485 nm. Curves represent mean ± SD (N = 3). For TEM images, scale bar, 250 nm

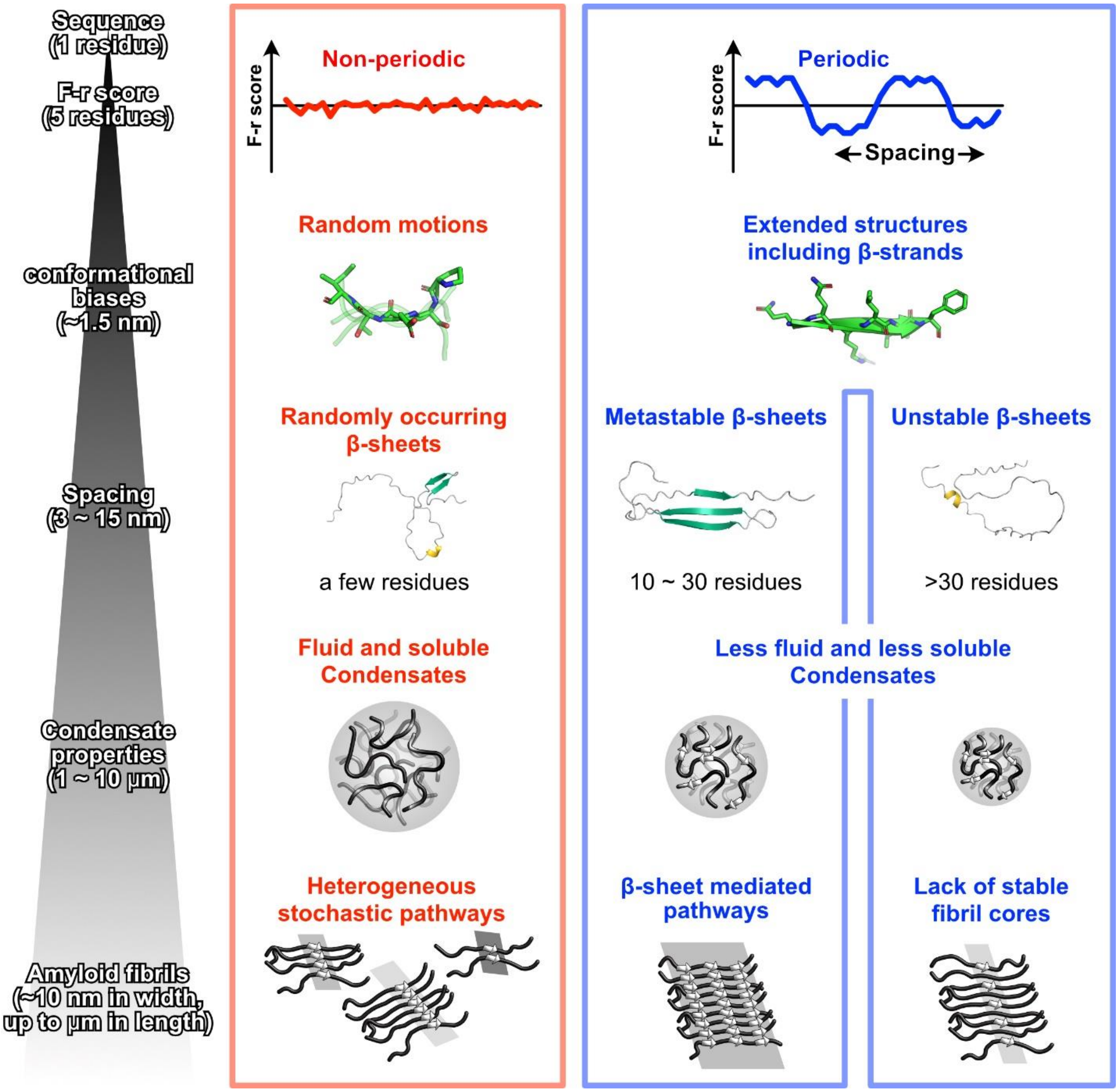


**Fig. S18. Proposed self-assembly mechanism for lag-series IDPs.**
The self-assembly of lag-series IDPs is proposed to be governed by the periodic arrangement of high and low F-r score regions, which encode local conformational biases. High F-r score segments tend to adopt extended structures, including β-sheets, promoting surface-to-surface interactions, whereas low F-r score segments favor random motions and point-to-point interactions. Short spacers (10–30 residues) between high F-r regions facilitate the formation of metastable β-sheets that can convert into intermolecular β-sheets and seed amyloid fibril growth. Longer spacers (>30 residues) keep high F-r regions apart, preventing stable β-sheet formation and favoring reversible condensates. In NP sequences, the randomized distribution of F-r scores leads to predominantly random motions, resulting in fluidic, highly dynamic condensates and heterogeneous amyloid formation pathways.

**SI References**


1. S. Seabold, J. Perktold, "Statsmodels: Econometric and Statistical Modeling with Python" in *Proceedings of the 9th Python in Science Conference* (2010; https://conference.scipy.org/proceedings/scipy2010/seabold.html), pp. 92–96.
2. N. Sekiyama, K. Takaba, S. Maki-Yonekura, K. I. Akagi, Y. Ohtani, K. Imamura, T. Terakawa, K. Yamashita, D. Inaoka, K. Yonekura, T. S. Kodama, H. Tochio, ALS mutations in the TIA-1 prion-like domain trigger highly condensed pathogenic structures. *Proc. Natl. Acad. Sci. U. S. A.* **119**, 1–12 (2022).
3. J. Schindelin, I. Arganda-Carreras, E. Frise, V. Kaynig, M. Longair, T. Pietzsch, S. Preibisch, C. Rueden, S. Saalfeld, B. Schmid, J. Y. Tinevez, D. J. White, V. Hartenstein, K. Eliceiri, P. Tomancak, A. Cardona, Fiji: An open-source platform for biological-image analysis. *Nat. Methods* **9**, 676–682 (2012).
4. D. A. Case, K. Belfon, I. Y. Ben-Shalom, S. R. Brozell, D. S. Cerutti, I. T.E. Cheatham, V. W. D. Cruzeiro, T. A. Darden, R. E. Duke, G. Giambasu, M. K. Gilson, H. Gohlke, A. W. Goetz, R. Harris, S. Izadi, S. A. Izmailov, K. Kasavajhala, A. Kovalenko, T. R. Krasny, T. Kurtzman, T. S. Lee, S. LeGrand, P. Li, C. Lin, J. Liu, T. Luchko, R. Luo, V. Man, K. M. Merz, Y. Miao, O. Mikhailovskii, G. Monard, H. Nguyen, A. Onufriev, F. Pan, S. Pantano, R. Qi, D. R. Roe, A. Roitberg, C. Sagui, S. Schott-Verdugo, J. Shen, C. L. Simmerling, N. R. Skrynnikov, J. Smith, J. Swails, R. C. Walker, J. Wang, L. Wilson, R. M. Wolf, X. Wu, Y. Xiong, Y. Xue, D. M. York, P. A. Kollman, *AMBER 2020* (University of California, San Francisco, 2020).
5. V. Hornak, R. Abel, A. Okur, B. Strockbine, A. Roitberg, C. Simmerling, Comparison of multiple amber force fields and development of improved protein backbone parameters. *Proteins Struct. Funct. Genet.* **65**, 712–725 (2006).
6. P. S. Shabane, S. Izadi, A. V. Onufriev, General Purpose Water Model Can Improve Atomistic Simulations of Intrinsically Disordered Proteins. *J. Chem. Theory Comput.* **15**, 2620–2634 (2019).
7. J. P. Ryckaert, G. Ciccotti, H. J. C. Berendsen, Numerical integration of the cartesian equations of motion of a system with constraints: molecular dynamics of n-alkanes. *J. Comput. Phys.* **23**, 327–341 (1977).
8. H. Mull, O. Beckstein, Technical Report : SPIDAL Summer REU 2018 Dihedral Analysis in MDAnalysis. **6**, 1–4 (2018).
9. N. Michaud-Agrawal, E. J. Denning, T. B. Woolf, O. Beckstein, MDAnalysis: A toolkit for the analysis of molecular dynamics simulations. *J. Comput. Chem.* **32**, 2319–2327 (2011).
10. R. Gowers, M. Linke, J. Barnoud, T. Reddy, M. Melo, S. Seyler, J. Domański, D. Dotson, S. Buchoux, I. Kenney, O. Beckstein, MDAnalysis: A Python Package for the Rapid Analysis of Molecular Dynamics Simulations. *Proc. 15th Python Sci. Conf.*, 98–105 (2016).
11. R. V. Mannige, J. Kundu, S. Whitelam, The Ramachandran Number: An Order Parameter for Protein Geometry. *PLoS One* **11**, 1–14 (2016).
12. N. Yoshida, The Reference Interaction Site Model Integrated Calculator (RISMiCal) program package for nano- and biomaterials design. *IOP Conf. Ser. Mater. Sci. Eng.* **773** (2020).

13. A. Kovalenko, F. Hirata, Potentials of mean force of simple ions in ambient aqueous solution. II. Solvation structure from the three-dimensional reference interaction site model approach, and comparison with simulations. *J. Chem. Phys.* **112**, 10403–10417 (2000).
14. W. L. Jorgensen, J. Chandrasekhar, J. D. Madura, R. W. Impey, M. L. Klein, Comparison of simple potential functions for simulating liquid water. *J. Chem. Phys.* **79**, 926–935 (1983).
15. F. Delaglio, S. Grzesiek, G. W. Vuister, G. Zhu, J. Pfeifer, A. Bax, NMRPipe: A multidimensional spectral processing system based on UNIX pipes. *J. Biomol. NMR* **6**, 277–293 (1995).
16. W. Lee, M. Tonelli, J. L. Markley, NMRFAM-SPARKY: Enhanced software for biomolecular NMR spectroscopy. *Bioinformatics* **31**, 1325–1327 (2015).
17. W. Lee, A. Bahrami, H. T. Dashti, H. R. Eghbalnia, M. Tonelli, W. M. Westler, J. L. Markley, I-PINE web server: an integrative probabilistic NMR assignment system for proteins. *J. Biomol. NMR* **73**, 213–222 (2019).
18. Y. Shen, A. Bax, Protein backbone and sidechain torsion angles predicted from NMR chemical shifts using artificial neural networks. *J. Biomol. NMR* **56**, 227–241 (2013).
19. A. Micsonai, F. Wien, É. Bulyáki, J. Kun, É. Moussong, Y. H. Lee, Y. Goto, M. Réfrégiers, J. Kardos, BeStSel: A web server for accurate protein secondary structure prediction and fold recognition from the circular dichroism spectra. *Nucleic Acids Res.* **46**, W315–W322 (2018).
20. J. Jumper, R. Evans, A. Pritzel, T. Green, M. Figurnov, O. Ronneberger, K. Tunyasuvunakool, R. Bates, A. Žídek, A. Potapenko, A. Bridgland, C. Meyer, S. A. A. Kohl, A. J. Ballard, A. Cowie, B. Romera-Paredes, S. Nikolov, R. Jain, J. Adler, T. Back, S. Petersen, D. Reiman, E. Clancy, M. Zielinski, M. Steinegger, M. Pacholska, T. Berghammer, S. Bodenstein, D. Silver, O. Vinyals, A. W. Senior, K. Kavukcuoglu, P. Kohli, D. Hassabis, Highly accurate protein structure prediction with AlphaFold. *Nature* **596**, 583–589 (2021).
21. W. Kabsch, C. Sander, Dictionary of protein secondary structure: Pattern recognition of hydrogen-bonded and geometrical features. *Biopolymers* **22**, 2577–2637 (1983).
22. Y. Kanda, Investigation of the freely available easy-to-use software 'EZR' for medical statistics. *Bone Marrow Transplant.* **48**, 452–458 (2013).
